# Striatal Pathways for Action Counting and Steering

**DOI:** 10.1101/2025.11.02.686102

**Authors:** Isabella P. Fallon, Marina Roshchina, Feiyang Hong, Sofia Fernandez, Shaolin Ruan, Henry H. Yin

**Author notes:** Correspondence to Henry Yin.

## Abstract

The basal ganglia (BG) are critical for coordinating voluntary movements, yet their precise contribution remains a subject of debate. Using a novel operant counting task, we trained mice to perform a specific number of lever presses to obtain a reward, enabling quantification of continuous kinematics, discrete actions, and action sequences. Optogenetic manipulations of direct pathway (dSPN) and indirect pathway (iSPN) striatal projection neurons exert bidirectional and dissociable influences on both movement steering and action count progression: activation of dSPNs extends press sequences as if resetting an internal accumulator, whereas activation of iSPNs prematurely terminates them, mimicking completion of the count. In vivo calcium imaging reveals distinct yet intermixed populations of dSPNs and iSPNs representing either lever approach or count progression, with ramping activity patterns consistent with accumulation and discharge dynamics. Importantly, the difference between dSPN and iSPN population activity scales with proximity to both spatial and count-based goals, unifying discrete and continuous control within a push–pull model. These findings establish the BG as a central circuit for integrating kinematic and sequential representations to monitor and steer progress toward behavioral goals.

## Introduction

The basal ganglia (BG) are a set of subcortical nuclei that have been implicated in many functions^1–3^. Although BG dysfunction is known to underlie the movement deficits of Parkinson’s disease and related disorders, the precise computations performed by BG circuits during normal behavior remain controversial. Classic theories emphasize the BG’s role in action selection. According to this view, the BG selects a single action through the direct pathway while suppressing competing actions via the indirect pathway, to enable discrete choice between behavioral options^2,4^. The details of each action (e.g., direction or speed) are assumed to be determined outside the BG. On the other hand, recent work suggests that the BG plays a role in top-down specification of continuous movement kinematics, such as velocity and position^5–7^.

These two views—categorical selection versus continuous control—have often been treated as mutually exclusive, yet it is possible that BG output may unify both processes by representation progress toward specific behavioral goals. During natural actions, animals must track their advancement through both physical space and internal task structure. They must monitor not only *where* they are moving but also how far along they are within a learned sequence. Indeed previous studies have shown that the dorsolateral striatum (DLS) contributes to both the initiation and execution of action sequences and to the continuous pursuit of spatial goals ^6,8,9^. Given these results, it is possible that BG circuits compute internal variables, such as elapsed time or number of completed actions, that help organisms determine when to transition from one phase of behavior to the next^10,11^.

A critical challenge in testing this idea is to design tasks that dissociate discrete action counts from continuous movement variables. Conventional behavioral tasks do not require the animal to estimate or monitor the number of actions. Consequently, neural activity measured during these tasks cannot distinguish between control of movement kinematics per se and representation of an internal count or goal progress. Addressing this limitation requires a behavioral paradigm in which we can simultaneously monitor and independently manipulate discrete action counts and continuous kinematic control.

Here, we developed a novel operant counting task in which mice must perform an exact number of lever presses before entering a reward port. This task allows us to simultaneously quantify discrete actions (e.g., individual lever presses), kinematics (e.g., position and velocity measures as mice move toward goal targets), and action sequences (e.g., a bout of lever pressing followed by reward port entry), while recording and manipulating neural activity in the striatum. Using pathway-specific optogenetics and calcium imaging, we found that direct and indirect pathways exert opposing yet complementary influences on both steering and counting. Moreover, calcium imaging revealed distinct, intermixed populations of striatal neurons representing either spatial approach or count accumulation, with ramping activity suggesting integrative computation over the sequence. The net difference between direct and indirect pathway activity tracked proximity to both spatial and count-based goals, supporting a push–pull model in which the BG regulate behavioral progress toward internal and external targets.

Together, these findings suggest that the BG are not limited to selecting discrete actions or specifying their kinematic parameters, but can also provide an internal measure of goal proximity that governs when to continue, reset, or terminate a behavioral sequence. This integrative perspective bridges the categorical action selection and continuous action specification views of BG function, and provides a mechanistic framework for understanding how complementary pathways in the BG steer organisms toward both physical and abstract goals.

## Results

### Mice learn to count lever presses

Nonverbal counting is a highly conserved process across many species, including mice^12–14^. We leveraged this ability by training mice to produce a specific number of lever presses to earn a reward. Food was not delivered if the mice failed to achieve the exact press count and break the infrared sensor in the reward port within 2 seconds (**Fig. 1, Extended Data Fig. 1**). In a subset of experiments, we also varied the count requirement (3, 5, and 7 presses), and the mice adjusted their behavior accordingly. The mean and standard deviation of the press count scaled proportionally with the count requirement (**Fig. 1e),** so counting behavior exhibits the scalar property, which has been previously shown for both counting and timing in animals^13–15^. In other words, count variability (standard deviation) increases proportionally with its mean. For the remainder of the experiments, mice were always trained on a count of 5.

**Fig. 1.**
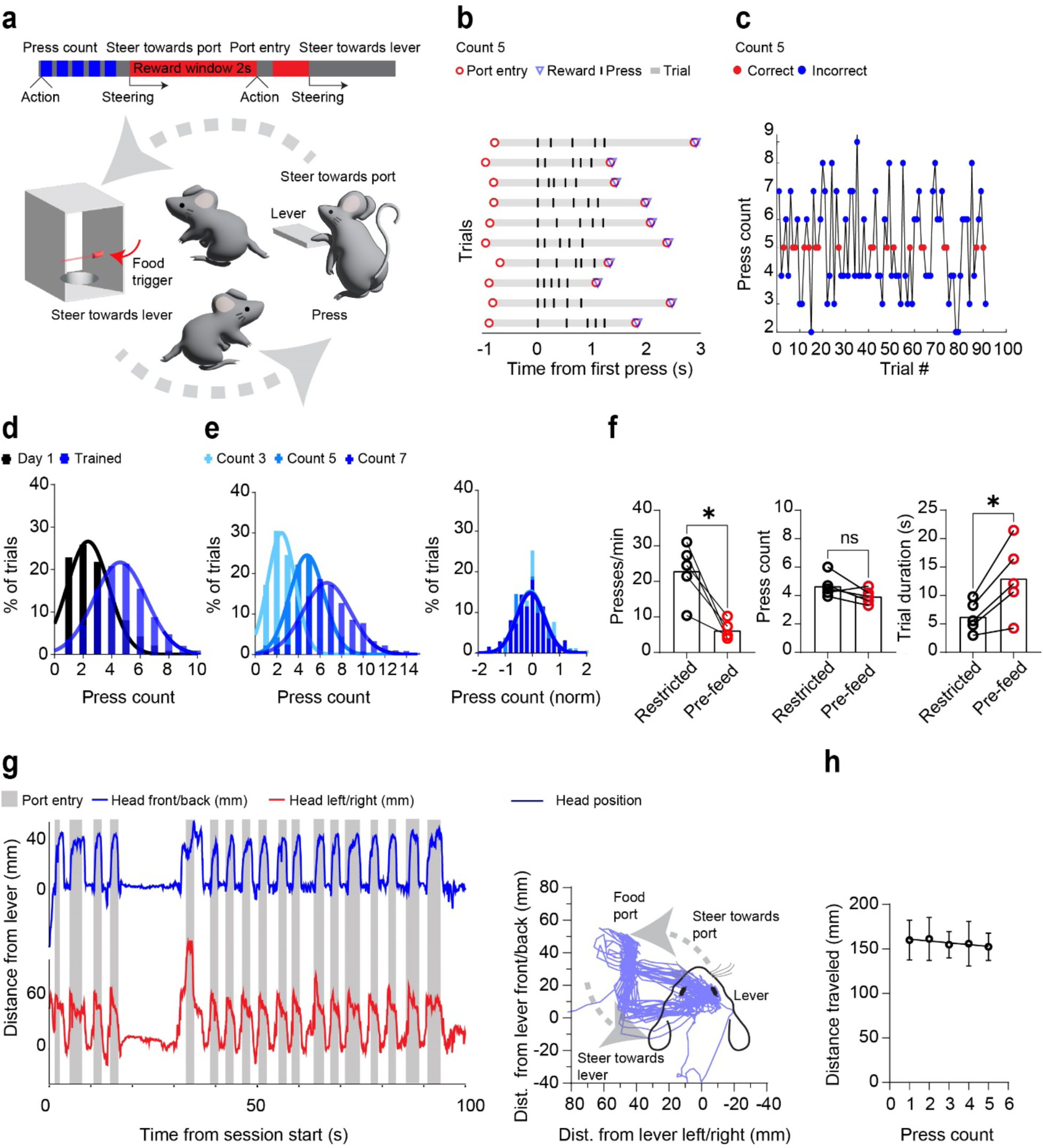
An operant counting task to allow simultaneous measurement of discrete action selection and continuous kinematics. **a)** When mice engaged in the press counting task, three phases were distinguishable: steering towards the lever, press counting, and steering towards the reward port. Reward was only delivered when mice entered the port after producing the required number of presses. **b)** A representative raster plot example of successful count-5 trials. Presses and port entries are aligned to the first press of the trial. **c)** A representative example of the number of presses in each consecutive trial for a trained mouse. **d)** Average press count increased during training. In trained mice (n = 30), the mean was similar to the count requirement. **e)** The distribution of the press counts for mice (n = 4) trained on a count of 3, 5, and 7 exhibited the scalar property: variability counting behavior scales proportionally with the mean of the count. **f)** The average press rate of pre-fed mice (n = 5) was significantly decreased (Paired t-test*, p =* 0.016). However, the average press count in each trial did not change following pre-feeding (Paired t-test, *p =* 0.1206). Average press trial durations were significantly increased by pre-feeding (Paired t-test, *p =* 0.0167). These observations suggest that mice did not use timing to decide when to enter the reward port. **g)** Representative trajectory of head position during the counting task. The mouse continuously transitioned between the lever and the port. **h)** The distance traveled did not correlate with the press count (n = 8, R^2^ = 0.002, *p =* 0.75). Bars represent the mean. Data points represent the mean ± SEM. Histogram bars represent the % of all trials across all mice. * *p <* 0.05.

It is important to note that the exact count requirement in our task is important as animals must rely on their internal count representation, rather than sensory feedback such as the sound of reward delivery. In contrast, mice trained on a conventional fixed-ratio (FR) task pressed in bouts, but the mean of their press bout distribution did not center around the ratio requirement (**Extended Data Fig. 2d**). This is because the FR schedule does not require a press bout for reward and simply delivers reward whenever the scheduled number of presses has been performed. Consequently, some mice use the strategy of pressing once or twice and checking the reward port instead of relying on a press bout to decide when to enter the reward port (**Extended Data Fig. 2**). Some could also rely on detecting the sound of reward delivery rather than monitoring the number of actions completed. FR schedules thus fail to dissociate motor execution from internal estimation processes, since reward predictability enables behavior driven by elapsed time or auditory cues rather than action count per se.

It is also possible that mice used time spent pressing rather than the number of presses to decide when to transition to the food port on our counting task. The BG have also been implicated in interval timing in the seconds to minutes range^16^, and recent work has suggested that animals use their own behavioral states and online perceptual feedback for timing^17,18^. According to one model, counting and timing share similar underlying mechanisms ^15^. If the decision to enter the port is based on timing only, then the mouse is expected to enter the reward port after a specific time interval. However, if the decision is based on counting, the mice should approach the reward port only after a certain number of presses, regardless of the time spent pressing. To determine whether mice relied on interval timing or press counting, we used devaluation by pre-feeding, which is expected to slow down goal-directed lever pressing^33^. Devaluation significantly decreased the press rate, significantly increasing the time spent completing the count, yet the count did not change significantly (**Fig. 1f**). This result confirmed that the performance was goal-directed, sensitive to outcome devaluation^19^, and that mice used press number rather than time to decide when to stop pressing and enter the reward port.

The lever and reward port represented the key spatial goals in this task, and trained mice transitioned between them on a consistent path (**Fig. 1g, h**). We included both male and female mice in our experiments and observed no significant differences in their task performance **(Extended Data Fig. 1f**).

### Direct and indirect pathways bidirectionally regulate steering and action counting

The striatum is the input nucleus of the BG and contains two classes of spiny projection neurons (SPNs) that bidirectionally control BG output^20–22^. We used excitatory and inhibitory opsins to selectively manipulate the activity of direct pathway projection neurons (dSPNs), which express D1-like dopamine receptors, and indirect pathway projection neurons (iSPNs), which express A2a adenosine receptors. For inhibition, the Cre-dependent inhibitory opsin (stGtACR2) was injected into the DLS of D1-cre or A2a-cre mice^23^. For excitation, the Cre-dependent excitatory opsin (ChR2) was injected (**Extended Data Fig. 3**)^24^.

In the first experiment, optogenetic stimulation was delivered immediately after the second press on 30% of runs and was delivered randomly to avoid anticipation. The hemisphere contralateral to the lever was chosen for testing first, since prior work has shown that unilateral striatal lesions and activation cause contraversive deficits^25–27^.

When we inhibited dSPNs unilaterally after the second press, mice usually steered ipsiversively towards the reward port (**Fig. 2a, Video 1**). In contrast, inhibition of iSPNs resulted in contraversive steering toward the lever (**Fig. 2b, Video 2**). Unilateral excitation of dSPNs produced contraversive steering, similar to inhibition of iSPNs (**Video 3**). Excitation of iSPNs, however, resulted in ipsiversive steering similar to dSPN inhibition (**Fig. 2c-d, Extended Data Fig. 4e-f, Video 4**). These findings revealed that selective activation of direct and indirect pathways could steer the mice in opposite directions toward the immediate goal targets (either the lever or reward port). On the other hand, inhibition of these pathways produced the opposite effects as those from excitation. These observations suggest that dSPNs and iSPNs are responsible for bidirectional control of steering direction.

**Fig. 2.**
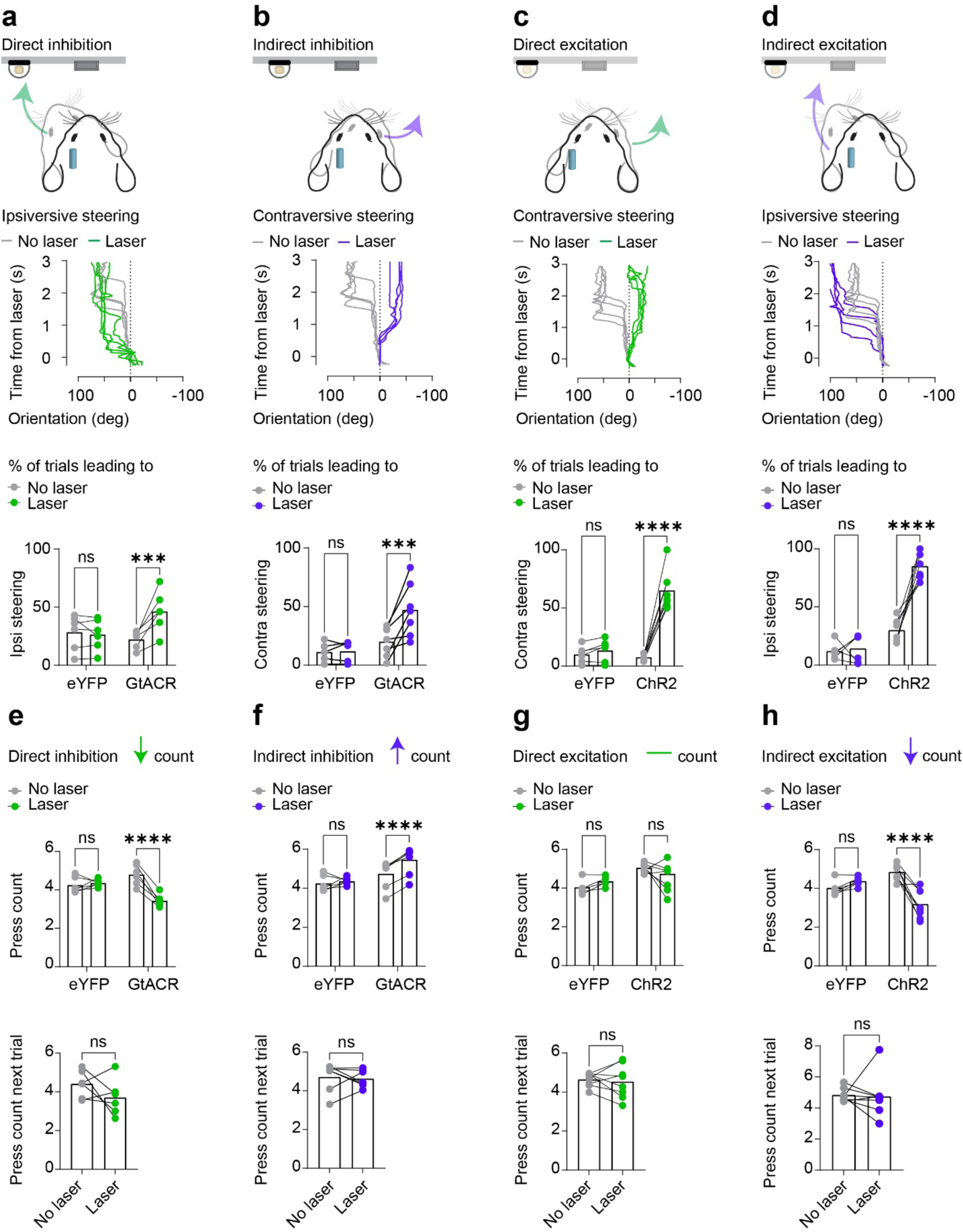
Bidirectional optogenetic manipulation reveals opposite and dissociable roles of direct and indirect pathways in counting and steering. **a-h)** For inhibition, D1-cre (n = 6) and A2a-cre (n = 7) mice were injected with a Cre-dependent inhibitory opsin SIO.GtACR2. For excitation, D1-cre (n = 8) and A2a-cre (n = 8) mice were injected with a Cre dependent excitatory opsin DIO.ChR2. Controls received injections eYFP (n = 6). Unilateral inhibition (500ms, 3-5mW, continuous) or excitation (500ms train duration, 10ms pulse width, 20-30hz, 5-7mW) of the hemisphere ipsilateral to the port was triggered by the second press. **a)** Direct pathway inhibition caused ipsiversive steering towards the reward port. The percentage of 2^nd^ presses that led to ipsiversive steering was increased by inhibition (Two-way ANOVA, significant interaction F (1,10) = 16.54, *p =* 0.002. Fisher’s LSD post hoc comparison revealed that inhibition of dSPNs led to more ipsiversive steering (*p <* 0.0007)). **b)** Indirect pathway inhibition caused contraversive steering towards the lever. The percentage of 2^nd^ presses that led to contraversive steering was increased by inhibition (Two-way ANOVA, significant interaction F (1,11) = 9.96, *p =* 0.0118. Fisher’s LSD post hoc comparison revealed that inhibition of iSPNs led to more contraversive steering (*p <* 0.0006)). **c)** Direct pathway excitation led to contraversive steering towards the lever. The percentage of 2^nd^ presses that led to contraversive steering was increased by excitation (Two-way ANOVA, significant interaction F (1,12) = 63.84, *p =* 0.0001. Fisher’s LSD post hoc comparison revealed that excitation of dSPNs led to more contraversive steering (*p <* 0.0001)). **d)** Indirect pathway excitation led to ipsiversive steering towards the port. The percentage of 2^nd^ presses that led to ipsiversive steering was increased by excitation (Two-way ANOVA, significant interaction F (1,12) = 28.98, *p =* 0.0002. Fisher’s LSD post hoc comparison revealed that excitation of iSPNs led to more ipsiversive steering (*p <* 0.0001)). **e)** Direct pathway inhibition reduced the number of presses (Two-way ANOVA, significant interaction F (1,10) = 33.38, *p =* 0.0002. Fisher’s LSD post hoc comparison revealed that inhibition of dSPNs reduced pressing (*p <* 0.0001)). On the next trial after stimulation, the press count was reset and was no different than the press counts without stimulation (Paired t-test, *p =* 0.230). **f)** Indirect pathway inhibition increased press count (Two-way ANOVA, significant interaction F (1,11) = 13.20, *p =* 0.00039. Fisher’s LSD post hoc comparison revealed that inhibition of iSPNs led to more pressing (*p <* 0.0001)). On the next trial after stimulation, the press count was reset and was no different than control trials (Paired t-test, *p =* 0.73). **g)** Direct pathway excitation did not change the number of presses (Two-way ANOVA, asignificant main effect of virus F (1, 12) = 12.31, *p =* 0.004, no significant main effect of laser stimulation, F (1,12) = 0.008, *p =* 0.92 and no significant interaction F (1,12) = 2.8, *p =* 0.12). On the next trial after stimulation, the press count remained unchanged (*p =* 0.95). h) Indirect pathway excitation reduced the number of presses (Two-way ANOVA, significant interaction F (1, 12) = 29.97, *p =* 0.0001. Fisher’s LSD post hoc comparison revealed that excitation of iSPNs reduced press count (*p <* 0.0001)). On the next trial after stimulation, the press count was reset and was no different than control trials (Paired t-test, *p =* 0.8529). Bars represent the mean. ** *p <* 0.01, *** *p <* 0.001, **** *p <* 0.0001.

Consistent with the effects on steering, dSPN inhibition and iSPN excitation had similar effects on counting, as measured by the number of lever presses on each trial. They reduced the press count and caused mice to steer toward the reward port (**Fig. 2e, h**). On the other hand, although dSPN excitation and iSPN inhibition both produced contraversive steering, they did not have the same effects on counting: dSPN excitation after press 2 did not change the count, whereas iSPN inhibition transiently increased the count, without affecting the count on the next trial (**Fig. 2f, g**). These results suggest that the effects on steering and counting could be partially dissociated.

### Count-dependent effect of stimulation

One possible explanation for these observations is that steering towards the reward port necessarily leads to count termination and reward port entry, and that steering the animal towards the lever necessarily leads to count increase. Although the effects on counting were generally consistent with the effects on steering, an exception was found with dSPN excitation, which did not increase the count as expected given the steering effect toward the lever side. If stimulation directing mice toward the reward port influenced only steering, the mouse is expected to resume pressing immediately after the stimulation ended, rather than entering the reward port. However, when stimulation steered the mice ipsiversively towards the reward port, they did not resume pressing after the end of stimulation but continued to move and entered the reward port, as if they had finished the count, since count termination required reward port entry (**Fig. 3a)**. On the next trial, they appeared to reset their press count to the learned requirement, starting the press count over (**Fig. 2e, h**).

**Fig. 3.**
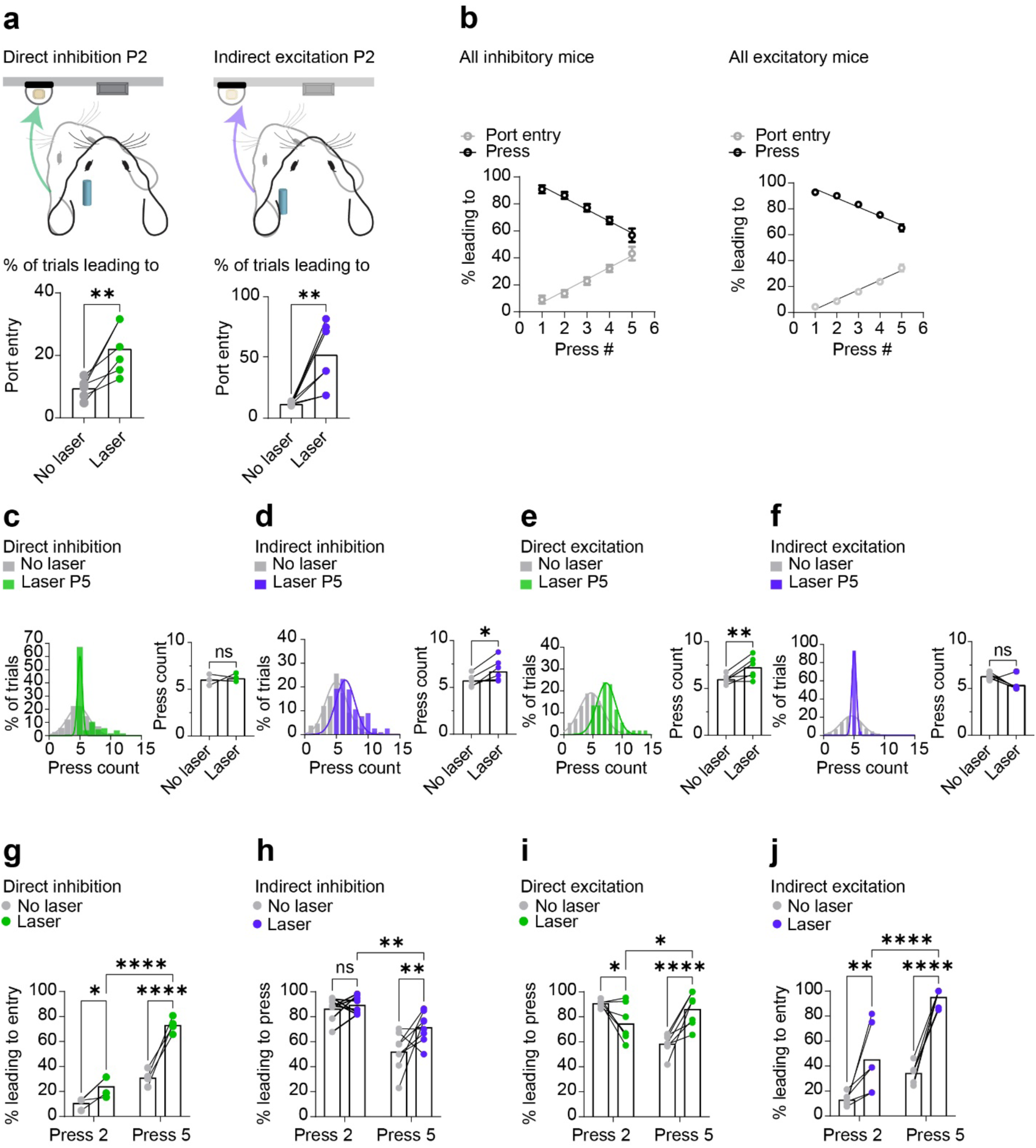
Stimulation effects on counting depend on the current count. **a-f)** Unilateral inhibition (500ms, 3-5mW, continuous) or excitation (500ms train duration, 10ms pulse width, 20-30hz, 5-7mW) of the hemisphere ipsilateral to the port was triggered by the 2^nd^ or 5^th^ press. **a)** Direct pathway inhibition and indirect pathway excitation both promoted reward port entry (direct pathway inhibition: paired t-test, *p =* 0.009, indirect pathway excitation: *p =* 0.0001**). b)** Without stimulation, the probability of a port entry increases with press number whereas the probability of another lever press decreased with press number (GtACR: n = 13; ChR2: n = 15). **c)** Direct pathway inhibition on the 5^th^ press narrowed the distribution of counts of 5 but did not change the mean press count compared to no laser trials with a minimum of 5 presses (Paired t-test, *p =* 0.74). **d)** Indirect pathway inhibition on the 5^th^ press caused a rightward shift in the distribution and significantly increased the press count compared to no laser trials with a minimum of 5 presses (Paired t-test, *p =* 0.016). **e)** Direct pathway excitation on the 5^th^ press caused a rightward shift in the distribution and significantly increased the press count compared to no laser trials with a minimum of 5 presses (Paired t-test, *p =* 0.008). **f)** Indirect pathway excitation on the 5^th^ press narrowed the distribution but did not significantly change the mean press count compared to no laser trials with a minimum of 5 presses (Paired t-test, *p =* 0.06). **g)** Direct pathway inhibition led to a greater probability of port entry when inhibition occurred on the 5^th^ press (Two-way ANOVA, significant interaction F (1,6) = 14.57, *p =* 0.008. Fisher’s LSD post hoc tests revealed that there was a greater increase in port entry when stimulation was delivered on the 5^th^ press (P2: *p <* 0.0124, P5: *p <* 0.0001). **h)** Indirect pathway inhibition increased pressing (Two-way ANOVA, significant interaction F (1,17) = 6.917, *p =* 0.0198). Fisher’s LSD post hoc tests revealed that only inhibition on the 5^th^ press increased pressing (P2: *p* > 0.05, P5: *p =* 0.0019, *p =* 0.0012). **i)** Direct pathway excitation increased pressing (Two-way ANOVA, significant interaction F (1,12) = 53.84, p = 0.0001. Fisher’s LSD post hoc tests revealed that excitation on the 5^th^ press led to a greater increase in pressing (P2: *p =* 0.01, P5: *p <* 0.0001). **j)** Indirect pathway excitation on the 5^th^ press increased port entry (Two-way ANOVA, significant interaction F (1,10) = 7.104, *p =* 0.0237. Fisher’s LSD post hoc tests revealed that excitation on the 5^th^ press increased port entry (P2: *p <* 0.0012, P5: *p <* 0.0001). Data represents mean ± SEM. * *p <* 0.05, ** *p <* 0.01, **** *p <* 0.0001.

We also compared the effects of stimulation after different press numbers. As mice approached the press count goal, the probability that each press would lead to a port entry increases, but the probability of performing another press decreases (**Fig. 3b).** If stimulation only produced a steering effect, then the timing of stimulation during the press run would not matter, i.e., it would produce the same effect regardless of the press count at which stimulation was delivered. We compared the effects of optogenetic stimulation early in the press bout (press 2) to those late in the press bout (press 5). If dSPN inhibition and iSPN excitation only steered mice toward the port, independently of the current count, the timing of the stimulation should not matter: stimulation would be expected to result in port approach with equal probability after press 2 or press 5. However, dSPN inhibition and iSPN excitation led to significantly more port entries when stimulated after the 5^th^ press than after the 2^nd^ press (**Fig. 3c, f, g, j, Extended Data Figs. 4a, d, 5a-b, g-h, Videos 5-6**). Thus, the effects on steering towards the reward port depended on the current count.

Since iSPN inhibition increased the total press count, we also evaluated whether the effects on pressing depended on press count. We found that inhibition of iSPNs at the end of the press bout led to more pressing than inhibition early in the press bout (**Fig. 3d, h, Extended Data Figs. 4b, 5c-d, Video 7**). Interestingly, excitation of dSPNs after press 5 also led to more pressing, as if the presses they had just performed were not sufficient to complete the count requirement. Excitation of dSPNs late in the press bout also led to more presses than activation early in the press bout (**Fig. 3e, i, Extended Data Figs. 4c, 5e-f, Video 8)**.

To test if the data was consistent with an internal count reset mechanism, we simulated count reset, and compared the simulation results to the experimental data. Simulation of count estimation with a 50% partial reset reproduced the behavioral effects of direct pathway activation: stimulation on either press 5 or press 2 made mice behave as though their internal counter had been partially reset, leading to extended press sequences (**Extended Data Fig. 6a-b).**

To summarize, activation of dSPNs resulted in more pressing, as if the count had been partially reset, while activation of iSPNs made the mice check the reward port, as if they had just completed the required press count. Although stimulation-induced steering toward the reward port led to early termination of the count, this effect was greater when stimulation was delivered after press 5 than after press 2. More importantly, while both dSPN activation and iSPN inhibition on press 2 caused contraversive steering, they had different effects on the press count. Only iSPN inhibition increased the count.

### Stimulation of the other hemisphere dissociates effects on steering and counting

We also stimulated the striatum in the other hemisphere contralateral to the port and on the side of the lever. In this hemisphere, dSPN and iSPN inhibition briefly paused pressing while mice remained oriented forward. But importantly this pause did not affect press count (**Extended Data Fig 7, Videos 9-12).** In addition, although excitation of dSPNs on press 2 often caused contraversive steering toward the port, it did not affect the average press count. Excitation after press 5, however, resulted in more pressing (**Extended Data Fig. 7c, g, Videos 13-14**). These results indicate that dSPN activation in either hemisphere resulted in longer press runs, even though stimulation in different hemispheres produced steering in opposite directions and toward different targets (lever vs. reward port). This is a clear dissociation between stimulation effects on steering and counting.

Conversely, activation of iSPNs after press 2 or press 5 resulted in ipsiversive steering toward the lever side. However, this did not result in lever pressing, but instead terminated the current press count, and often resulted in a complete 360-degree turn to the reward port. This increased port entries (**Extended Data Fig. 7d, h, Videos 15-16**). These results demonstrate that even though iSPN activation in different hemispheres produced steering in opposite directions, it reliably terminated the press count by steering mice to the reward port.

Optogenetic manipulation of the other hemisphere confirmed that the direction of steering is independent of the effect on counting. dSPN activation prolonged the ongoing progression toward the count goal, as if they had not yet completed the task. In contrast, iSPN activation had the opposite effect: it terminated the current count, and mice behaved as if they were finished. Thus, the direct and indirect pathways have opposing and dissociable effects, not only on steering toward spatial goal targets but also on steering toward an action sequence count goal.

### Distinct striatal cell populations represent target steering and count accumulation

Our optogenetic results suggested that the outputs from the direct and indirect pathways can influence two distinct features of behavior on this task: (1) steering, which can be measured by continuous kinematic variables and (2) a press count sequence, which involves discrete presses. One possibility is that striatal neurons regulate both steering toward a target and lever pressing, consistent with a role in organizing action sequences. Another possibility is that distinct populations of SPNs regulate steering and pressing. To evaluate these two possibilities, we recorded neural activity from dSPNs and iSPNs using *in-vivo* calcium imaging while mice performed the press count task (**Fig. 4, Extended Data Fig. 3).**

**Fig. 4.**
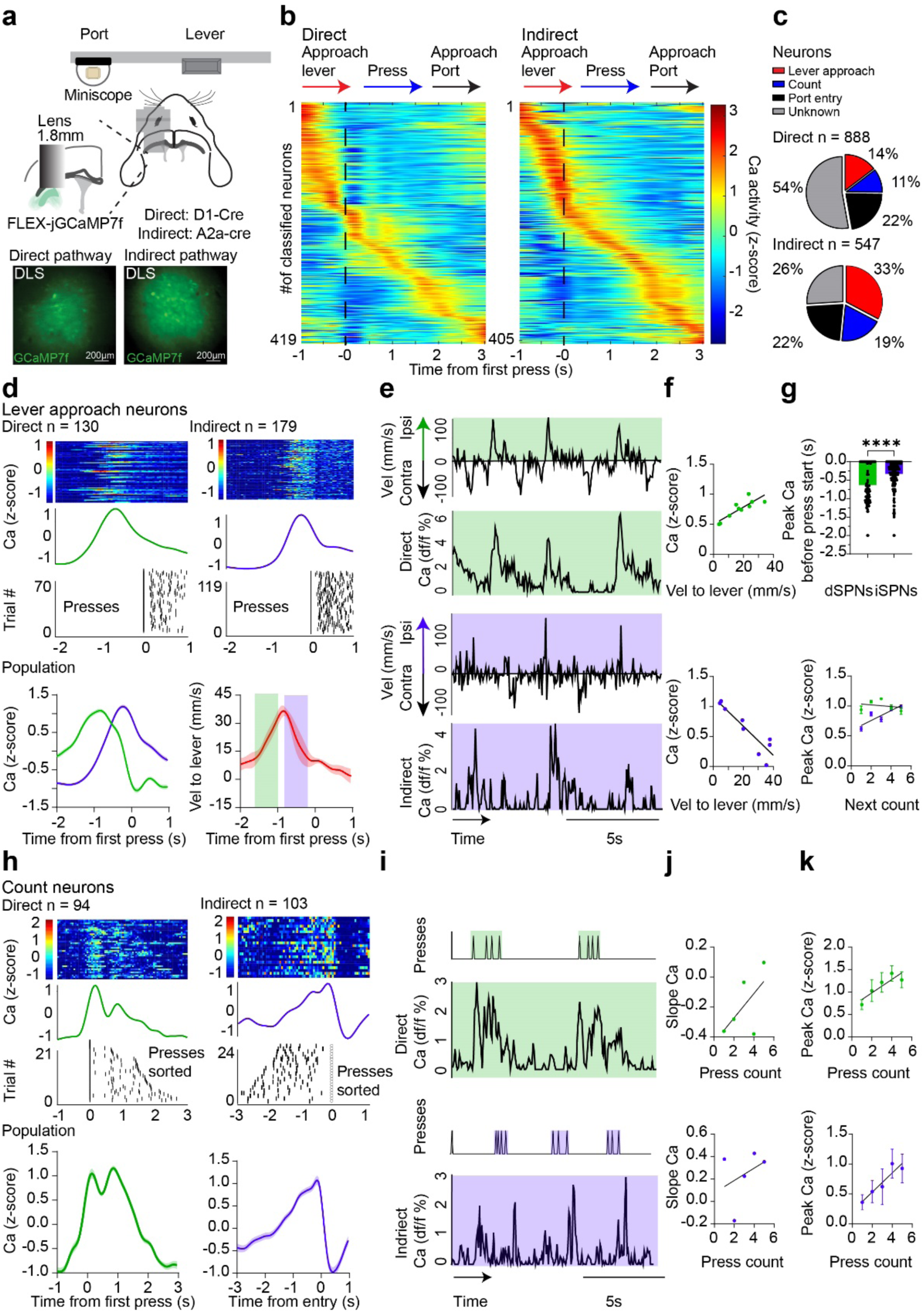
Distinct striatal cell populations mediate goal directed steering and action sequence counting behavior. **a)** In-vivo calcium imaging of dSPNs and iSPNs. The hemisphere ipsilateral to the reward port was recorded (A2a: n = 5, D1: n = 4). Representative examples of calcium signals in D1+ and A2a + mice. **b)** The population of task-related direct and indirect pathway neurons increased activity at distinct components of the press count behavior and were classified by their temporal relationship to behavior. **c)** The proportion of neurons in each group. **d)** (Top) Representative examples of direct and indirect pathway lever approach neurons aligned to press run start. (Bottom) the population of lever approach neurons increased activity as mice contraversively approached the lever. The shaded bars show the time window used for computing correlation between neural activity and behavior. **e)** A representative direct pathway lever approach neuron and the mice’s velocity. A representative indirect pathway lever approach neuron and the mice’s velocity. **f)** Direct pathway lever approach neuron population activity was positively correlated with velocity toward the lever (R^2^ = 0.73, *p =* 0.0016). Indirect pathway lever approach neuron population activity was negatively correlated with velocity toward the lever (R^2^ = 0.63, *p =* 0.005). **g)** For lever approach neurons, dSPN population activity peaked before iSPN population activity (Unpaired t-test, *p <* 0.0001). The peak activity of dSPN and iSPN lever approach neurons was not correlated with the next count (dSPNs: R^2^ = 0.06, *p =* 0.688; iSPNs: R^2^ = 0.76, *p =* 0.051). **h)** (Top) Representative examples of direct and indirect pathway count neurons aligned to press run start. (Bottom) The population of direct and indirect pathway count neurons increased activity during pressing. **i)** A representative direct pathway count neuron and the mice’s presses. A representative indirect pathway count neuron and the mice’s presses. **j)** The slope of direct pathway count neuron activity was not correlated with count (R^2^ = 0.37, *p =* 0.27). The slope of indirect pathway count neuron activity was not correlated with count (R^2^ = 0.12, *p =* 0.55). **k)** The peak of direct pathway count neuron activity at the beginning of the count was positively correlated with count (R^2^ = 0.77, *p =* 0.04). The peak of indirect pathway count neuron activity at the end of the count (entry at run end) was positively correlated with count (R^2^ = 0.87, *p =* 0.01). Data represents mean ± SEM.

We identified two major populations of neurons based on their relationship with task-relevant variables. The first population is activated during mice’s approach toward the lever. Both dSPNs and iSPNs were found in this “lever approach” population (**Fig. 4b-d**). The dSPNs were activated earlier, corresponding to a positive correlation with lever-directed velocity, whereas iSPNs were activated later. Despite being active before pressing, the activity of lever approach neurons was not related to the next press count (**Fig. 4e-g**). When mice steered in the other direction, towards the port, iSPN activity was higher than dSPN activity (**Extended data Fig. 8c**).

The second population of neurons (“count neurons”) was activated during the press count and steering towards the reward port (**Fig. 4b, c, h**). This population also included both dSPNs and iSPNs, but they showed different patterns. While dSPNs decreased activity during the press run, iSPNs increased activity. The peak but not the slope of activity in dSPN and iSPN count neuron populations covaries with press count (**Fig. 4i-k**). In addition, because dSPN count neurons peaked before iSPN count neurons, they were negatively correlated with velocity towards to port whereas iSPN count neurons were positively correlated with velocity towards the port (**Extended data Fig. 8a-b**). These results suggest that the count-related neurons are not just related to pressing per se; rather their activity appear to reflect accumulation and discharge in an integrator^13,15^. This accumulation process dictates when to move ipsiversively to the port.

Some neurons showed activity related to reward port entry (**Fig. 4b-c** and **Extended Fig. 9**). This “port neuron” population was not clearly related to leftward and rightward steering kinematics. These neurons were not modulated by reward delivery, and neither were the other two groups of neurons (**Extended Data Fig. 8d-f and Extended Data** Figs. 9 **c-f**). As we could not collect detailed kinematic data once the mice were in the reward port, it is unclear what type of behavioral variables are represented by this population.

When the average activity of all recorded neurons was used in the analysis, there was no clear difference between dSPN and iSPN activity during lever and port approach (**Extended Data Fig. 10i-k**). Finally, to determine whether the different neuronal populations were spatially segregated, we also analyzed the locations of neurons recorded during calcium imaging. Different populations were spatially intermixed, showing no clear segregation (**Fig. 5**).

**Fig. 5.**
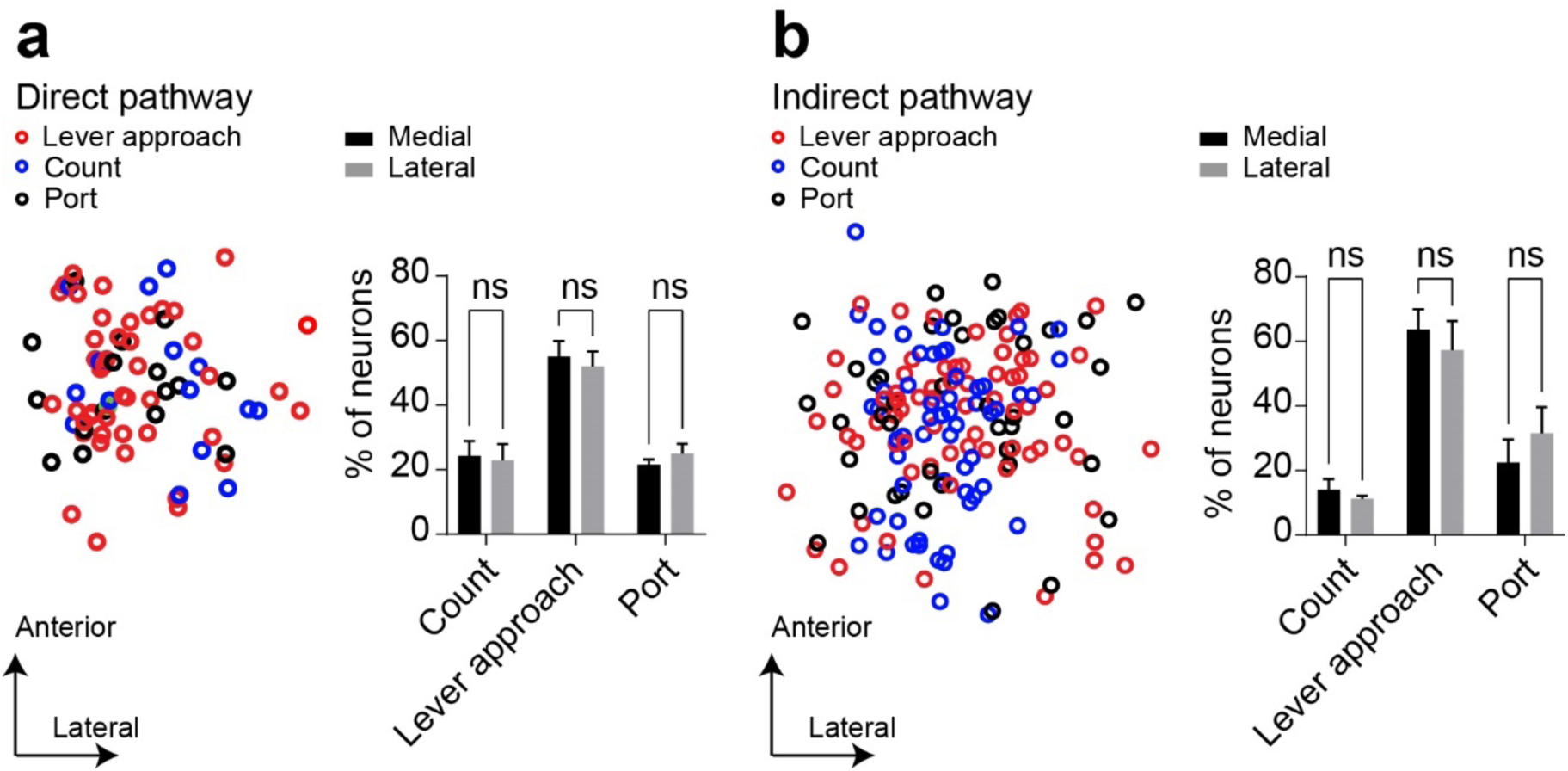
Different functional classes of SPNs are spatially intermixed. **a)** Representative spatial positions of dSPNs from different functionally defined populations. The proportion of neurons in each population did not depend on the spatial position (Two-way ANOVA, significant main effect of cell group (F (2, 25) = 36.63, *p =* 0.0001), no significant main effect of spatial position (F (1, 24) = 0.01, *p =* 0.92), and no interaction (F (2, 24) = 0.34, *p =* 0.70). **b)** Representative spatial positions of iSPNs from different functionally defined populations. (Two-way ANOVA, significant main effect of cell group (F (2, 18) = 28.83, *p <* 0.0001), no significant main effect of spatial position (F (1, 18) = 9.950e-008, *p =* 0.99) or interaction (F (2, 18) = 0.76, *p =* 0.47). Data represents mean ± SEM.

### Difference between dSPN and iSPN activity represents proximity to goal

It is well-established that dSPNs inhibit BG output nuclei such as the substantia nigra pars reticulata (SNr), releasing inhibition on downstream structures, whereas the iSPNs promote SNr inhibition (**Fig. 6a**)^4,28^. The anatomical organization suggests that the net output of this circuit may be proportional to the difference between dSPN and iSPN outputs. We used calcium imaging to record activity in these two pathways in the same mice and analyzed the relationship between net BG output (dSPN activity minus iSPN activity) and behavior on this task. We replicated the results of our previous recordings and found both dSPNs and iSPNs in lever approach and count neuron populations (**Fig. 6**). During approach, net BG output scaled with how close the mouse was to the lever, indicating that these circuits monitor goal proximity rather than movement velocity (**Fig. 6b-c, Extended Data Fig. 10a, c).** On the other hand, in the count population, the net output correlated with reward port proximity and not velocity (**Fig. 6d-e, Extended Data Fig. 10b,d**). When the net activity of all populations combined was analyzed, the relationship was no longer observed (**Extended Data Fig. 10e-h**). These results suggest that, on the counting task, net BG outputs from different functional modules represent proximity to physical goal targets and count completion. These could represent error signals for different controlled variables.

**Fig. 6.**
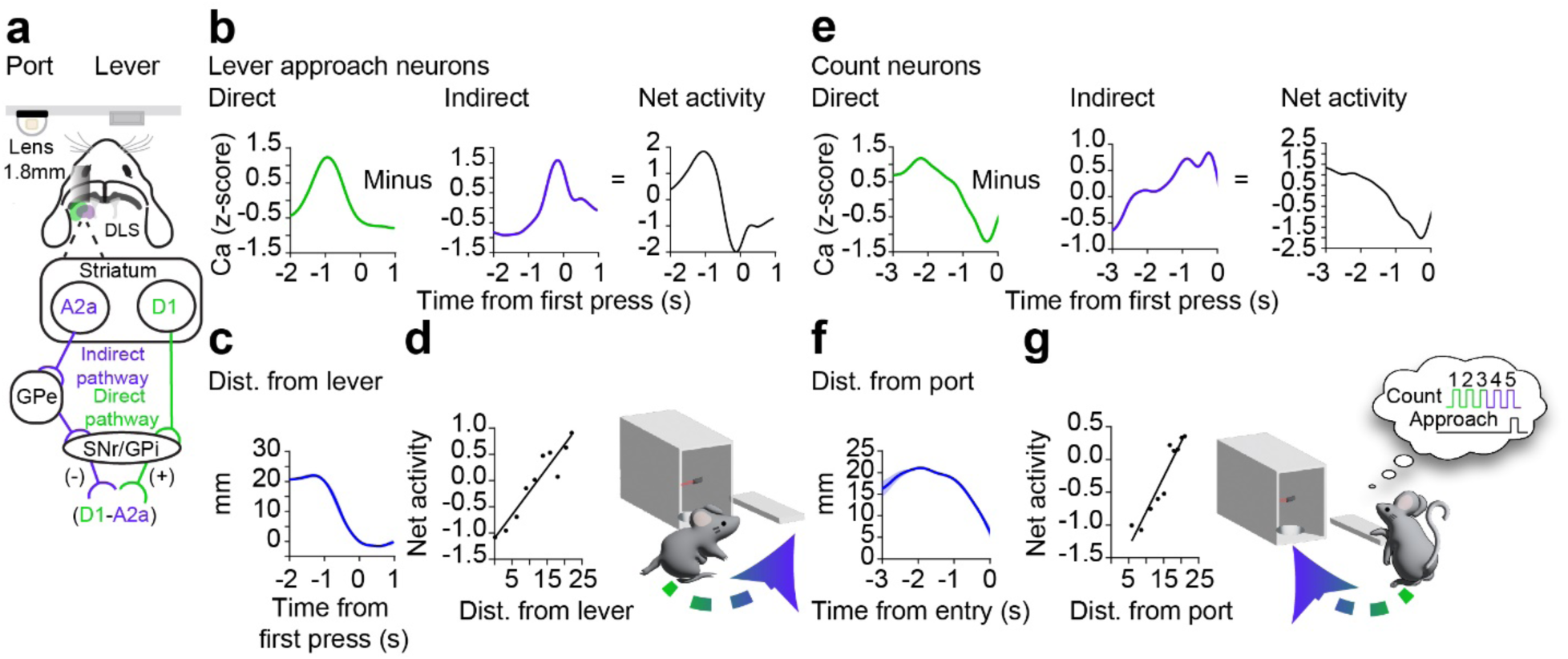
The difference between direct and indirect pathway activity determines goal proximity. **a)** For downstream neurons receiving basal ganglia output, the net input is roughly proportional to subtracting iSPN activity from dSPN activity. To record direct and indirect pathway activity in the same animal, D1-cre or A2a-cre mice were co-injected with a Cre-ON calcium indicator and a Cre-OFF calcium indicator (n = 4). **b)** Representative example of direct, indirect, and net lever-approach neural activity as the mouse approached the lever. **c)** Representative example of the same mouse’s average distance from the lever during the same time window as in (b). **d)** There was a positive correlation between the average net activity of lever approach neurons and distance from the lever. (R^2^ = 0.89, *p <* 0.0001). **e)** Representative example of a mouse’s direct, indirect, and net count neuron activity as the mouse approached the port. **f)** Representative example of the same mouse’s average distance from port during the same time window as in (e). **g)** There was a positive correlation between the average net activity of count neurons and the mice’s average horizontal distance from the port. (R^2^ = 0.90, *p <* 0.0001). Data represents mean ± SEM.

## Discussion

We found that direct and indirect pathway neurons contribute to both the accumulation of discrete actions (lever press count) and the specification of continuous movement parameters (steering toward spatial targets). Optogenetic manipulations showed that these pathways bidirectionally regulate both lever press count and steering behavior. Calcium imaging identified separate striatal populations for these functions, with their net output predicting proximity to spatial and count-based goals. Interestingly, none of the neuronal populations examined were directly related to reward, as reward delivery had no significant impact on neural activity in the DLS, in agreement with previous work ^6,9^. Moreover, selective optogenetic manipulations of dSPNs and iSPNs systematically biased steering behavior towards the lever and the reward port (**Figs. 2-3**). But the stimulation effects were not limited to steering. Activation of dSPNs increased the count (i.e. presses produced in a bout before reward port entry), whereas activation of iSPNs completed the count by causing a reward port entry.

One alternative explanation for the present results is that stimulation of dSPNs and iSPNs just causes contraversive and ipsiversive steering, and changes in action counting are a result of stimulation-induced steering. However, results from the optogenetic manipulation in the other hemisphere revealed that this explanation is inadequate (**Extended Data Fig. 7).** Direct pathway activation in the other hemisphere led to steering towards the reward port, but still had the same effect of increasing the count. Indirect pathway activation in the other hemisphere led to steering towards the lever, but had the same effect of reducing the count. In addition, while both activation of dSPNs and inhibition of iSPNs caused contraversive steering towards the lever, only inhibition of iSPNs changed the press count. The stimulation effects on steering can be dissociated from the effects on counting; changes in kinematics alone (such as direction of approach behavior) are not sufficient to cause changes in counting (**Fig. 3**).

Our calcium imaging results are consistent with optogenetic stimulation results. We found distinct striatal cell populations that mediate behavioral trajectories towards goal targets. These populations are selectively activated during different phases of the behavioral cycle: one was activated during lever approach, another during counting and approach to the reward port, and the last after mice entered the reward port. Activity of count-related dSPNs was correlated with the number of presses and exhibited a ramping-down pattern. This activity profile suggests that activating these neurons could reset the accumulator and increase the number of presses within a trial, an effect we indeed observed (**Fig. 3e**). In contrast, count-related iSPNs showed a ramping-up pattern that peaked as mice stopped pressing and entered the reward port. The activity of these neurons was positively correlated with press count, suggesting that activating these neurons may prematurely terminate the counting process, a prediction also supported by our data (**Fig. 2h**). The opposing ramping profiles of dSPNs and iSPNs resemble integration and discharge dynamics of an internal accumulator that tracks progress through an action sequence. Interestingly, this pattern is also consistent with previous models of interval timing and evidence accumulation^15,29^.

In addition, count neurons were also active during transition to the reward port. Termination of the count is indicated by reward port approach, and count neurons (both dSPNs and iSPNs) were related to velocity towards the port. dSPNs were negatively correlated with whereas iSPNs were positively correlated with velocity towards the port (**Extended Data Fig. 8**). On the other hand, lever approach dSPNs were positively correlated with velocity towards the lever (contraversive) and iSPNs showed the opposite relationship (**Fig. 4**).

Together, these findings suggest that dSPN and iSPN activity bidirectionally regulates some accumulation process in an internal integrator that regulates the number of actions generated within a learned sequence. Consistent with pace-making accumulator models, dSPNs act to fill the integrator, while iSPNs act to discharge it.^13,15,30^ Unlike simple accumulator models with static leak and time constants, however, the BG circuits appear to be characterized by the capacity for dynamic gating, which allows continuation and termination of sequences based on a learned criterion such as the press count.

Previous work has found SPN activity increases at the start or stop of actions, supporting discrete action selection or sequence control^31^ while others found continuous relationships to kinematics^6,32,33^. In agreement with other recent studies^9,34^, we found that striatal activity represented more than just start and stop of actions, but showed time varying relationship with kinematic variables. Our observations are consistent with the idea that striatal neurons are essential for both initiating actions and specifying action parameters^35^.

Some limitations of our study should be noted. First, because we were unable to track mouse behavior within the reward port, we could not fully characterize neuronal activity during this phase. Notably, neurons that increased activity after port entry were not related to reward delivery (**Extended Data Fig. 8**) and may instead reflect reward consumption or forward/backward approach behavior while in the port^9^. Future studies incorporating detailed monitoring of kinematics within the reward port will be necessary to determine the functional role of these neurons. We did not observe medial to lateral differences in the distribution of neuronal groups or significant variation in the effects of our optogenetic manipulations. This could result from cortical damage caused by the GRIN lens, which produced extensive damage to the sensorimotor cortex above the dorsal striatum. Thus, we cannot rule out the possibility that different neural dynamics might emerge with these cortical areas intact. It is also possible that our placement does not allow us to examine differences between DMS and DLS effectively. Future studies with more precise targeting will be needed to compare these two regions in counting and steering behavior.

Although both dSPNs and iSPNs are activated during different phases of the behavior on this task, dSPN activity consistently leads iSPN activity for both lever approach and count neurons, in agreement with previous work showing distinct patterns in dSPNs and iSPNs^11,36,37^. Previous work arguing for concurrent activity was based on photometry recordings with poor temporal and spatial resolution^38^. It is possible that striatal photometry recordings may reflect non-somatic calcium signals rather than somatic spiking-related calcium changes^39^. Our results show direct pathway activation typically leads indirect pathway activation, though the lag is variable. It remains unclear exactly how this lag is determined, whether by the timing of cortical or thalamic inputs to these populations or by intrinsic BG circuit properties.

Since dSPNs and iSPNs have opposite effects on BG output nuclei (e.g. from the SNr)^40^, it is possible that the relative activation of these two populations may determine net BG output at any time^41^. To test this hypothesis, we computed the difference between the population activity from dSPNs and iSPNs and analyzed the relationship between net BG output and behavior in the same animal. We found that the net output of lever approach and count neurons represents proximity to the lever and the reward port, respectively **(Fig. 6).** This result suggests a push-pull model in which the relative activation of these pathways determines the behavioral trajectory. The difference between the two populations may determine when count is terminated and reward port approach is initiated. Indeed, previous work has shown that activation of dopamine neurons, which modulate SPN activity, can increase movement velocity in reward approach behavior^42,43^. Because dopamine increases dSPN activity and reduces iSPN activity, it may increase the net difference between SPN modules and thus increase approach towards the goal target. Thus striatal dopamine level may bias the balance of dSPN/iSPN activity, dynamically regulating proximity representations in approach behavior.

According to one proposal for how BG circuits can implement action selection, the direct pathway selects the desired action, whereas the indirect pathway suppresses competing actions^2^. However, recent studies have shown the limitations of this model, demonstrating that the BG also play a critical role in specifying continuous movement kinematics^1,5,6,9,30^. Our findings show that both dSPNs and iSPNs can represent kinematics during goal-directed behavior, and that activation of these neurons can generate bidirectional steering. They suggest that, in thinking about BG function, it is important to move beyond the traditional conceptual framework that considers behavior in all-or-none, categorical terms. Rather the direct and indirect pathways jointly regulate an internal accumulator, which can be modeled with a leaky integrator. In this model, dSPNs provides the main inflow through disinhibition, and iSPNs providing leak or discharge. This model is supported by opposite ramping patterns of dSPNs and iSPNs during account, and by partial reset of the count after dSPN activation, and by premature termination of the count after iSPN activation. A similar architecture in a nested hierarchical organization can be used for integration at different time scales for behavioral units of different sizes. In this way, the BG can flexibly govern both discrete action counts and continuous kinematic parameters through dynamic, push–pull control rather than categorical selection alone.

In summary, our findings bridge the long-standing divide between BG models of action selection and kinematic control. They suggest that discrete and continuous control variables can both be controlled through neural dynamics in direct and indirect pathways that track behavioral progress. Distinct yet intermixed populations of dSPNs and iSPNs guide movements toward different goal types, whether a spatial target like a lever or reward port, or an abstract, sequential target, such as completing a prescribed number of actions. The absence of clear anatomical segregation in the functional populations of dSPNs and iSPNs suggests that these functional ensembles lack fixed molecular identity but rather emerge through instrumental learning. By uniting discrete action counting with continuous movement steering under a shared computational mechanism, our results reveal a unifying principle in which BG circuits monitor and regulate ongoing progress toward both physical and abstract goals.

## Acknowledgments

The authors would like to thank Fengxia Allen, Konstantin Bakhurin, Alexander Friedman, Jiwon Kim, and Guozhong Yu for their technical assistance, and Russell Ravenel for his valuable feedback on the manuscript. HHY is supported by DA040701, NS121253, and NS094754.

## Author contributions

I.P.F. and H.H.Y. designed experiments; I.P.F., M.R. and S.R. conducted optogenetic experiments; I.P.F. conducted behavioral experiments; I.P.F. and F.H. conducted calcium imaging experiments; I.P.F. and S.R. analyzed data; I.P.F. and H. H. Y. wrote the manuscript.

## Inclusion and Ethics

All animal experiments were conducted in compliance with the ethical guidelines approved by the Institutional Animal Care and Use Committee (IACUC) of Duke University. Both male and female mice were included in the study, and sex differences were evaluated. Animals were randomly assigned to experimental groups to minimize bias. Efforts were made to minimize discomfort.

## Data availability

The datasets generated and analyzed during the current study, along with the custom codes used for data analysis, are available from the corresponding author upon reasonable request.

## Methods

### Mice husbandry

To selectively target the direct and indirect pathways, D1-Cre (Drd1-Cre, 37156, The Jackson Laboratory, Maine, USA) and A2a-Cre (*Adora2a^tm1Dyj^*, 010687, The Jackson Laboratory, Maine, USA) aged 3-8 months were used. For control experiments, wild type (WT) (C57BL/6J, 000664, The Jackson Laboratory, Maine, USA) aged 3-4 months were used. All experimental procedures were pre-approved by the Animal Care and Use Committee at Duke University. All mice were grouped-housed, 2-5 mice per cage, and kept on a 12:12hr light: dark cycle. Testing occurred during the light phase. Outside of operant experimentation, mice had ad libitum access to food and water.

### Viral constructs

For inhibition experiments, the Cre dependent pAAV1_hSyn1-SIO-stGtACR2-FusionRed was used (105677, Addgene, MA, USA). For control experiments rAAV5-hSYN-eYFP was used (UNC Core, NC, USA). For excitation experiments, pAAV5/EF1a-DIO-hChR2-EYFP(E123T/T159C)-eYFP was used (35509, Addgene, MA, USA). For single-color calcium imaging experiments, pGP-AAV9-syn-Flex-jGCaMP7f-WPRE (104492, Addgene, MA, USA) was used. For two-color calcium imaging experiments, AAV-hSyn-DO(FAS)-jRCaMP1b-WPRE (20689, Duke Core, NC, USA) and pGP-AAV9-hSyn-F-jGCaMP7f-WPRE (104492, Addgene, MA, USA) or pAAV.Syn.Flex.NES-jRCaMP1b.WPRE.SV40 (100850, Addgene, MA, USA) and AAV9-CBA-DO(FAS)-GCaMP6s (pBK1557, Duke Core, NC, USA).

### Surgery

Mice were anesthetized in chamber filled with 3% isoflurane before being placed into a stereotaxic frame (David Kopf Instruments, Tujunga, CA) and were maintained at 1.0-1.5% during surgery.

For optogenetic experiments, 200nL of AAV-SIO-GtACR or 200nL of AAV-DIO-hChR2 was infused bilaterally into the DLS (A/P: +0.4 mm, M/L: +/-2.4 mm, D/V: -3.2 and -2.8 mm from skull surface) of A2a-Cre and D1-cre mice using a micro infusion pump (Nanoject 3000, Drummond Scientific). For optogenetic control experiments 200nL of AAV-eYFP was injected into WT mice. To allow diffusion of the virus, the microinjector remained in place for 4mins following infusion. Custom fabricated optic fibers (3mm below the ferrule, >85% transmittance, 105 μm core diameter) were implanted above the DLS (A/P: +0.4 mm, M/L: +/- 2.4, D/V: -2.6 mm from skull surface).

For single color calcium imaging experiments, 500nL of AAV-FLEX-GCaMP7f was infused into the DLS (A/P: +0.2 mm, M/L: + or - 2.2 mm, D/V: -2.8 to -2.0 mm, in 0.2 mm steps) of A2a-Cre and D1-Cre mice. The injection hemisphere was evenly split among groups. Mice were trained so that the recording hemisphere was always ipsilateral to the reward port.

For two color calcium imaging experiments, the same coordinates as single-color calcium imaging experiments were used. D1-cre (n=2) mice were co-infused with 500nL AAV-SYN-DO(FAS)-jRCaMP1b and AAV-SYN-FLEX-GCaMP7f (2:1) into the right hemisphere. A2a-Cre (n=2) mice were co-infused with 500nL AAV-SYN-FLEX-jRCaMP1b and AAV9-CBA-DO(FAS)-GCaMP6s (2:1) into the left hemisphere. This strategy ensured that the recording hemisphere was split evenly between mice and A2a+ cells always appeared red while D1+ cells always appeared green. Again, mice were trained so that the recording hemisphere was ipsilateral to the reward port.

For all calcium imaging experiments, cortical tissue was aspirated using iced PBS prior to the implantation (lens center A/P: +0.4 mm, M/L: +2.2 mm, D/V: -2.0 mm) of a GRIN lens (1.8 mm × 4.2 mm, Edmund Optics). A baseplate to attach the miniscope was affixed to the top of the skull using dental cement. All animals received post-operative care for 5 days following surgery and were given 1 week to recover before experimentation.

### Histology

Following the completion of experiments, mice were anesthetized with 3% isoflurane and transcardially perfused with 0.9% saline followed by 4% paraformaldehyde (PFA). Brains were extracted and kept in 4% PFA at 4°C for 24 hours. Brains were then transferred to a 30% sucrose solution and kept at 4°C until they sank to the bottom of the tube. For tissue sectioning, brains were immersed in Tissue-Tek O.C.T. and frozen at -80°C. Brain tissue was sectioned using a cryostat at 30 µm, placed directly on Superfrost Plus slides, and mounted with DAPI Vectashield (Vector Laboratories, CA). The entire striatum was sectioned to assess surgical placement and viral expression. Images were acquired using a LSM 710 confocal microscope (Zeiss, Oberkochen, Germany) at 20-40X objectives and processed using FIJI software.

### Behavior

Prior to operant experimentation, mice were food restricted and maintained at 85-90% of their body weight. For training, each mouse was placed into a soundproof operant chamber equipped with two levers on either side of a food magazine port containing an infrared beam inside (Med Associates Inc, VT, USA). At the start of each session, the house light was turned on and one lever was inserted. Mice were trained to obtain a food pellet reward (Bio-Serv 14mg Dustless Precision Pellets, Bio-Serv, NJ, USA) by pressing the lever. The relationship between lever pressing and reward was controlled by a custom Med-PC-IV program. Above each box was a blackfly camera to record continuous behavior during the session (BFS-U3-04S2M-CS, Teledyne FLIR, Virginia, USA). At the end of each session, the house light was turned off and the lever was retracted. The end of each session was triggered when mice reached 76 pellet rewards or exceeded 60 minutes, whichever occurred first.

### Fixed-ratio behavior

In a separate experiment mice (n = 11) were trained on a fixed ratio (FR) task. Mice were first trained on a fixed-ratio 1 (FR1) lever press schedule where they received a reward for each lever press, for 3 sessions (1 session/day). For the next 10-17 days, mice were trained to press the lever 5 times for a reward (FR5) until thy achieved all the required rewards (76) in the session before the time of the session was completed (60mins) for 3 consecutive days. For the next 13-15 days, mice were trained to press the lever 10 times for a reward (FR10) until they earned all the rewards in the session before the time of the session was completed (60mins) for 3 consecutive days. For the next 10-12 days, mice were trained to press the lever 15 times for a reward (FR15) until they achieved all the required rewards (76) in the session before the time of the session was completed (60mins) for 3 consecutive days.

### Action counting task

For the counting task, mice were first trained on a fixed-ratio 1 (FR1) lever press schedule in which they received a reward for each lever press, for 3 sessions (1 session/day). For the next 3 days, mice were trained to press the lever 5 times for a reward (FR5) until the mean of their press count distribution reached 2. In subsequent sessions, they were trained on count schedules, in which to receive a reward the animal must produce both the exact press count and reward port entry within 2 seconds of the last press. Reward port entry was signaled by breaking an infrared beam in the magazine port. The press counter was reset if the mice pressed more than 5, entered the port before reaching the correct number of presses, or if the mice waited longer than 2 s after the last press to enter the port. Mice were trained on count-3 schedule (3-4 days) until the mean of the press count distribution reached 3. They were then trained on a count-5 training schedule until their press count distribution remained stable (5±1) for 3 consecutive days. To test the scalar property of counting behavior, a new cohort of mice were also trained on a count-7 schedule until the mean of the press count distribution remained stable at 7 for 3 days.

In devaluation experiments, the same mice trained in the scalar experiments with the addition of one new mouse, were also given a devaluation test when trained on a count of 5. First, a baseline session was recorded in the fully trained mice. Then, for the experimental session the following day, mice were pre-fed 1.5g of pellets before being placed in the operant box. They were placed in the box immediately after eating all the pellets.

### Closed loop optogenetics

For optogenetic experimentation, mice were trained on the count 5 schedule as described above. A 470 nm DPSS laser (Shanghai Laser & Optics, Shanghai, China) was placed above each operant box and a 1.5 m sheathed fiber (105-µm core diameter, 0.22NA) was extended into the chamber and attached to the mouse’s fiber optic implant using a ceramic sleeve (Precision Fiber Produces, CA, USA). For inhibition experiments, the laser power was 3.5-5 mW. For excitation experiments, the laser power was 5-7 mW and was delivered at 20-30hz (10ms pulse width, 10 or 15 pulses). Stimulation duration was designed to cover one inter-press interval. In each optogenetic testing session, the laser was triggered immediately following 2 presses or 5 presses.. Photo-stimulation took place on 30% probability of trials. Stimulation frequency was controlled through an Arduino (Arduino, MA, USA). A Blackfly camera was affixed above the box to record behavior (BFS-U3-04S2M-CS, Teledyne FLIR, Virginia, USA) and was recorded at 50fps.

### In vivo calcium imaging

For calcium imaging experiments, mice were trained on a count-5 schedule (see behavior). Based on the optogenetic results, mice were trained such that the lever was contralateral to the recording hemisphere.

For single-color calcium imaging experiments (Fig. 4), a V3 UCLA blue LED miniscope (CA, USA) was affixed to the skull baseplate to record D1+ or A2a+ neurons during the counting task (see surgery for viral strategy). Calcium was imaged at 30 fps with 5-15% LED power. A Blackfly camera was affixed above the box to record behavior (BFS-U3-04S2M-CS, Teledyne FLIR, Virginia, USA) and was recorded at 100 fps. The miniscope camera and the behavior camera were triggered 5 minutes after the MED-PC-IV program was started using an Arduino, to allow mice to acclimate and begin pressing before recording. Once the cameras were triggered, the recording lasted 10 minutes. One recording was taken for each mouse.

For two-color calcium imaging experiments (Fig. 6) D1+ and A2a+ neurons were recorded in separate sessions. First, a V3 UCLA green LED miniscope (CA, USA) was affixed to the skull baseplate to record A2a+ neurons (see surgery for viral strategy) during the counting task. A2a+ neurons were imaged at 20 fps with 15-20% LED power. A Blackfly camera was affixed above the box to record behavior (BFS-U3-04S2M-CS, Teledyne FLIR, Virginia, USA) and was recorded at 100 fps. The miniscope camera and the behavior camera were triggered simultaneously with the MED-PC-IV program using an Arduino and recorded for 10 minutes. Second, a V3 UCLA blue LED miniscope (CA, USA) was affixed to the same mouse’s skull baseplate to allow recording of D1+ neurons (see surgery for viral strategy) during the counting task. D1+ neurons were imaged at 30 fps with 5-15% LED power. A Blackfly camera was affixed above the box to record behavior (BFS-U3-04S2M-CS, Teledyne FLIR, Virginia, USA) and was recorded at 100fps. The miniscope camera and the behavior camera were triggered simultaneously with the MED-PC-IV program using an Arduino and recorded for another 10 minutes. One session for each color was recorded for each mouse.

### Analysis procedures

To ensure proper alignment of data, all timestamps were collected using a Blackrock data acquisition system (NeuroPort System, Blackrock Microsystems). All data analyses were performed using custom scripts in MATLAB (MathWorks), NeuroExplorer 5.414 (Nex Technologies), Python, ImageJ (NIH), and Prism (GraphPad).

### Analyses of behavioral training data

During behavioral training sessions, event timestamps (lever presses, port entries, and pellet delivery) were extracted from Blackrock recordings using a custom MATLAB script and imported into NeuroExplorer for visualization and analysis. Press count distributions were created by summing up the number of presses between port entries and plotting a histogram of the press counts during the session. Press count distributions were monitored daily. Graphs and statistical analysis were performed in Prism and MATLAB; see captions for statistical details.

### Analyses of optogenetic data

For optogenetic experiments, continuous mouse positions were labeled and tracked using DeepLabCut (DLC) (https://deeplabcut.github.io/DeepLabCut/README.html)^44^. Video frame times, behavioral positions, and event timestamps (lever presses, port entries, and reward delivery) were extracted from Blackrock recordings using a custom MATLAB script and imported into NeuroExplorer for visualization and analysis. Body orientation was calculated by first drawing a line in the X-Y plane with the tail marker and the head marker when the mouse was facing the lever. Whether a trial led to steering was determined by deviation from the starting body orientation and the delay between the press and the next event (another press or entry). If there was a deviation in the starting body orientation but the delay did not exceed the mode value of their inter-press interval histogram, then the behavior was not considered. These classifications were corroborated by reviewing the videos. To calculate the press count for trials that contained stimulation, an interval was created between the first press before stimulation to the next port entry. The number of presses within that interval was quantified. These counts were compared to the average number of presses from no-stimulation trials in the previous session that had at least the same number of presses as the trial with photo-stimulation. For % leading to press or % leading to entry, the proportion of stimulation or control trials where a press at position *n* was followed by a press at *n+1* within a pre-specified time window was quantified. For Fig. 3 and Extended Data Figs. 5, 7, laser distributions reflect the distribution of press counts in stimulation trials.

### Simulation of count reset

We conducted a Monte Carlo simulation (1,000 trials) to investigate how internal count estimation is affected by memory resets at fixed points. In each trial, a target number of 5 presses was set. Internal estimates were modeled as Gaussian-distributed random variables with noise proportional to the target (standard deviation σ = 0.3 × target).

Three conditions were simulated:

1. **No Reset**: A single estimate was drawn from a normal distribution with increased variability (σ × 1.5), simulating degraded memory accuracy.
2. **Reset at 5 Presses**: The count was split into a fixed pre-reset value of 5 and a post-reset estimate drawn from N(μ = 0.5 × target, σ), reflecting a partial reset.
3. **Reset at 2 Presses**: Similar to the 5-press reset condition, but with the partial reset occurring earlier (pre-reset = 2).

Negative estimates were clipped to zero, and final counts were summed for reset conditions. Distributions of estimated total presses were compared using normalized histograms.

### Analyses of calcium imaging data

Mouse positions from the Blackfly video recordings were labeled and tracked using DeepLabCut^44^. Calcium recordings were motion corrected, denoised, and deconvolved using the analysis software Minian (https://github.com/denisecailab/minian)^45^. Miniscope frame times, video frame times, behavioral positions, calcium signals, and event timestamps (lever presses, port entries and rewards) were extracted from Blackrock recordings using a custom script and imported into NeuroExplorer for visualization and analysis. Graphs and statistical analysis were performed in Prism and MATLAB; see captions for statistical details.

### Classification

Neurons were classified according to the temporal distribution and occurrence of their activity peaks in relation to defined behavioral epochs, including the initiation and termination of the press sequence and entry into the reward port. Neurons were classified as “lever approach neurons” if the average activity across trials was above 1 standard deviation within 1.5-0 seconds before the start of the press run compared to activity during pressing. Neurons were classified as “count neurons” if the average activity across trials increased above 1 standard deviation during the press bout compared to activity before pressing. Neurons were classified as “port neurons” if the average activity over trials increased above 1 standard deviation within 500-750ms following port entry compared to activity during pressing.

To plot peri-event calcium activity, each cell was aligned to the start of the press run or entry at run end, binned in 50ms intervals, smoothed using a Gaussian filter (3 bins), and z-scored. For lever approach neuron correlations, behavior (left/right approach velocity) and activity were aligned to the start of the press run, extended to 2 secs before the run to capture lever approach velocity (contraversive) and 1 second after to capture the start of the press run. Each cell was binned in 50ms intervals and averaged across trials before being normalized using a vector-wise z-score. The correlation between neural activity and lever approach velocity was calculated across 10 bins during movement transition. For direct pathway count neuron correlations, all neurons were aligned to the start of different press counts (1, 2, 3, 4, and 5), a window was extended 0.5 seconds before press run start and 2 seconds after, binned in 50ms second intervals and normalized using a vector-wise z-score over the whole-time interval. The peak of the population between the start of the press run to 1 second after was recorded for each mouse and each count. To estimate the slope of the population ramp, we fit a linear regression line to activity aligned to different counts, starting at the beginning of the press run and extending 1.5 seconds afterward. For indirect pathway count neuron correlations, all neurons were aligned to the start of entry at run end following different press counts (1, 2, 3, 4, and 5), a window was extended 2 seconds before entry at run end and 1 second after, binned in 50ms intervals and normalized using a vector-wise z-score over the whole time interval. The peak of the population between 1 second before entry at run end to entry at run end was recorded for each mouse and each count. To estimate the slope of the population ramp, a line was fit to the activity aligned to different counts, starting from entry at the end of the press run and extending 1.5 seconds earlier.

### Spatial position

To analyze whether the proportions of each cell group varied based on their spatial position (DMS or DLS), a custom MATLAB script was used to plot cell groups on top of a maximum projection of the recording area. Then, an analysis window was drawn over the recording window and split in half to distinguish the DMS from the DLS. The proportion of cells in each group was then quantified.

### Estimating net activity

To obtain the net activity (**Fig. 6**), we used data from the two-color calcium imaging experiments with dSPN and iSPN data from the same animal (n = 4 mice), 2 mice from single-color direct pathway (D1+) recordings, paired with 2 mice from indirect pathway (A2a+) recordings with similar behavior. Neurons in each recording were first classified in the same way as described above. Next, all lever approach neurons in the D1+ and A2a+ recordings were aligned to the start of the press run with a window extended 2 seconds before to obtain activity during lever approach and 1 second after. All count neurons were aligned to entry at run end with a window extended 3 seconds before to obtain activity during pressing and port approach. The data was binned (50ms), normalized using the vector-wise z-score over the whole-time interval, and averaged across trials and then neurons. The net activity was computed by subtracting the mean A2a+ signal from the D1+ signal. This process was repeated for each mouse with 2-color imaging and each pair of mice with 1-color imaging. For **Fig.6 b, e** the data portrayed is smoothed using a gaussian filter (3bins). To correlate activity with distance from the lever or port, each mouse’s distance measure was aligned, binned and averaged exactly the same as above but not smoothed. Then the correlation between average activity and distance was taken for each mouse across 10 bins during movement transition.

