## Supplementary data for "Striatal Pathways for Action Counting and Steering"

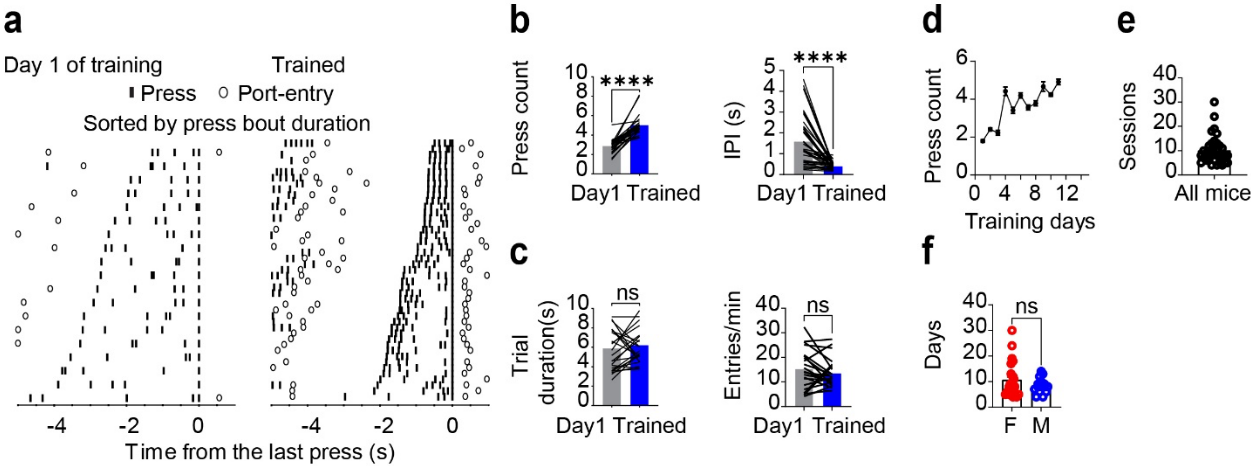


**Extended Data Fig. 1. Performance on the count task improves with training. a)** Representative raster plot examples of successful count-5 trials on the first day of training and the last day of training. Presses and port entries are aligned to the last press of correct trials (5$^{th}$ press). **b)** The mean press count in each trial increased after training (Paired t-test, $p < 0.0001$). The median inter-press interval decreased over training (Paired t-test, $p < 0.0001$). **c)** Mice maintained similar trial durations between day 1 and the final day of training (Paired t-test, $p = 0.8516$) because the rate of port-entry did not change over training (Paired t-test, $p = 0.296$). **d)** Representative example mouse showing the mean press count across training days. **e)** On average, mice took ~ 10 days to become proficient on the press count task. **f)** Males and females did not differ in the number of training days it took to become proficient on the counting task (Unpaired t-test, $p = 0.24$). Bars represent mean ± SEM. Bars represent mean. **** $p < 0.0001$.

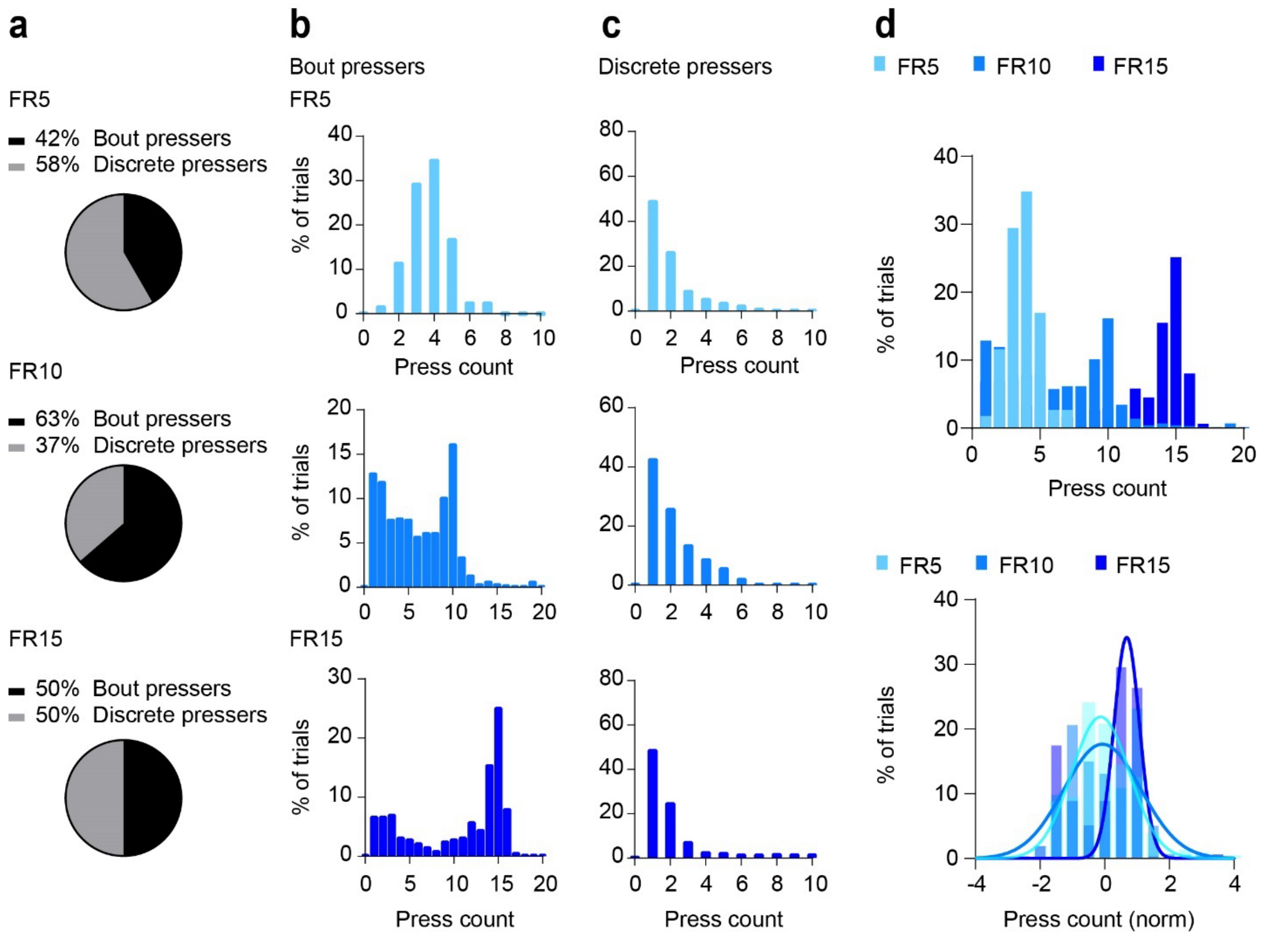


**Extended Data Fig. 2. Fixed ratio schedules do not reliably produce count behavior. a)** When trained on the fixed-ratio (FR) task, a proportion of mice did not learn to press in bouts (FR5 = 5 presses, FR10 = 10 presses, FR15 = 15 presses). **b)** Distribution of press counts. **c)** For mice that did not learn to press in bouts (discrete pressers), the press count distribution did not center around the press requirement. **d)** Press count distribution (all trials from all mice) did not overlap when press count is normalized, i.e. did not exhibit the scalar property. Thus, FR schedules are not very effective in training counting behavior in mice.

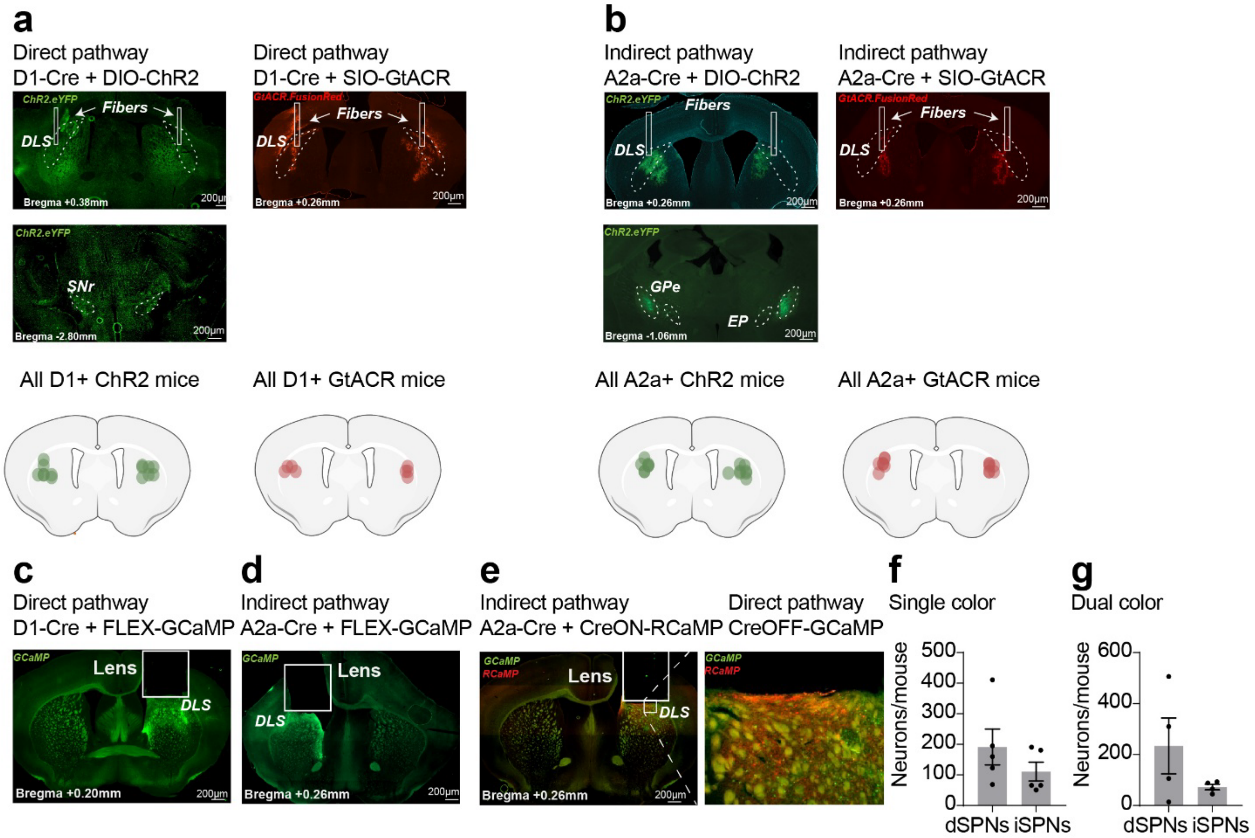


**Extended Data Fig. 3. Virus was selectively expressed in dSPNs and iSPNs. a)** Representative photomicrograph examples of DIO.ChR2 and SIO.GtACR2 opsins expressed in D1+ neurons in the striatum. Axons terminated in the SNr. **b)** Representative photomicrograph examples of DIO.ChR2 and SIO.GtACR2 opsins expressed in A2a+ neurons. Axons terminated in the GPe. Green and red dots on summary slices represent the estimated fiber tip locations from all mice. **c)** Example of FLEX.GCaMP7f expressed in D1+ neurons. **d)** Example of FLEX.GCaMP7f expressed in A2a+ neurons. **e)** Example of FLEX-RCaMP expressed in A2a+ neurons and DO(FAS)-GCaMP6s expressed in putative D1+ neurons. **f)** The number of neurons recorded in each animal, in single color experiments. **g)** The number of neurons recorded in each animal, in dual color experiments. Data represents mean ± SEM.

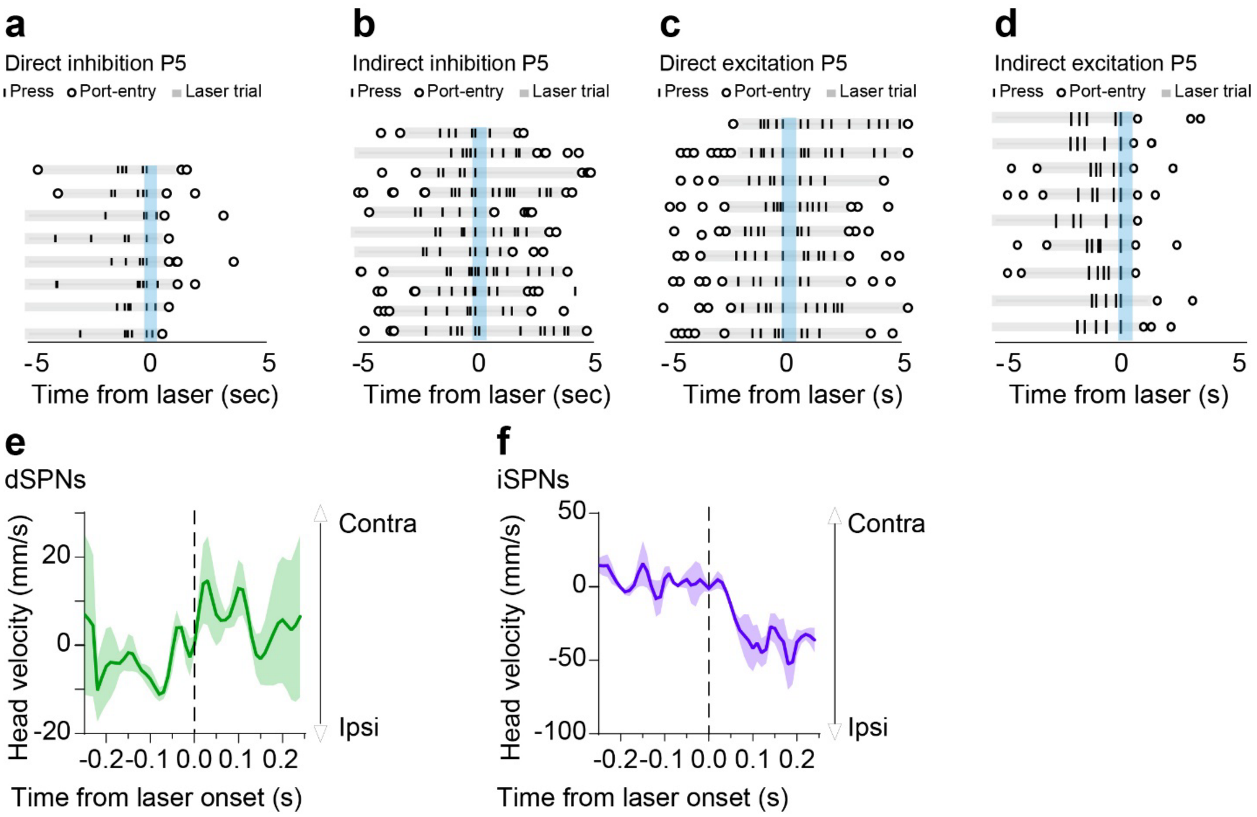


**Extended Data Fig. 4. Optogenetic manipulations of the direct and indirect pathways have opposite effects on pressing and head velocity. a-d)** Unilateral inhibition (500ms, 3-5mW) or excitation (500ms, 20-30Hz, 5-7mW) of the hemisphere ipsilateral to the port was triggered by the fifth press. Blue bar indicates timing of stimulation. **a)** Representative raster plot of direct pathway inhibition, which led to port entry. **b)** Representative raster plot of indirect pathway inhibition, which led to more pressing. **c)** Representative raster plot example of direct pathway excitation, which led to more pressing. **d)** Representative raster plot of indirect pathway excitation, which led to port entry**. e)** Excitation of dSPNs increased contraversive velocity of the head. **f)** Excitation of iSPNs increased ipsiversive velocity of the head. Data represent mean ± SEM.

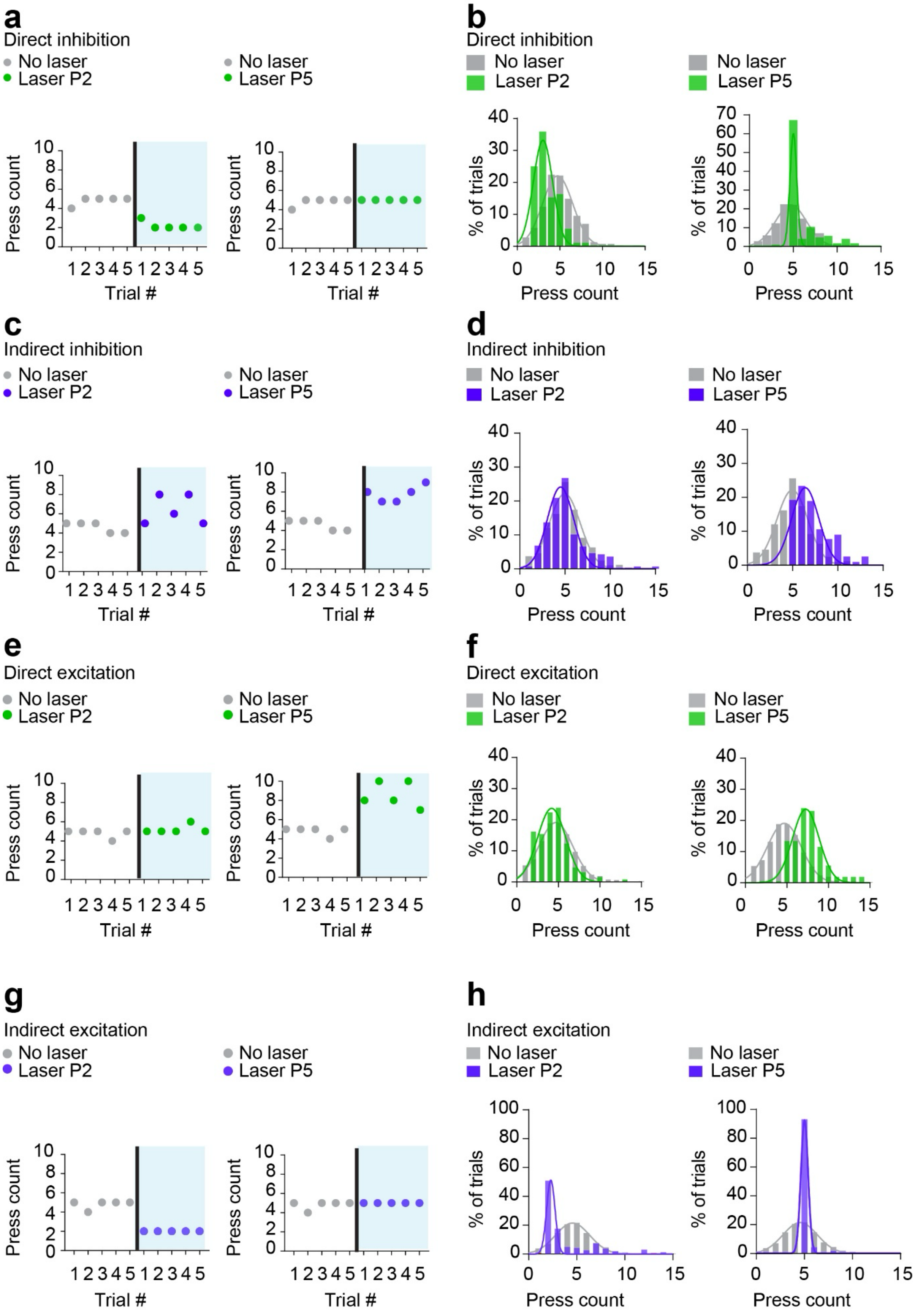

a
Direct inhibition
No laser
Laser P2
No laser
Laser P5
Press count
Trial #
b
Direct inhibition
No laser
Laser P2
No laser
Laser P5
% of trials
Press count
c
Indirect inhibition
No laser
Laser P2
No laser
Laser P5
Press count
Trial #
d
Indirect inhibition
No laser
Laser P2
No laser
Laser P5
% of trials
Press count
e
Direct excitation
No laser
Laser P2
No laser
Laser P5
Press count
Trial #
f
Direct excitation
No laser
Laser P2
No laser
Laser P5
% of trials
Press count
g
Indirect excitation
No laser
Laser P2
No laser
Laser P5
Press count
Trial #
h
Indirect excitation
No laser
Laser P2
No laser
Laser P5
% of trials
Press count

**Extended Data Fig. 5. Individual examples and population distributions of optogenetic effects on count. a-h)** Unilateral inhibition (500ms, 3-5mW) or excitation (500ms, 20-30Hz, 5-7mW) of the hemisphere ipsilateral to the port was triggered by the $2^{nd}$ or $5^{th}$ press. **a)** Representative examples of press counts from non-consecutive, selected trials with no stimulation, or with direct pathway inhibition triggered on press 2 or press 5. **b)** For all mice, direct pathway inhibition on press 2 caused a leftward shift in the press count distribution and on press 5, caused a narrowing of the distribution to counts of 5. **c)** Representative examples of a mouse's press counts from non-consecutive, selected trials with no stimulation, or with indirect pathway inhibition triggered on press 2 or press 5. **d)** For all mice, indirect pathway inhibition on press 2 did not change the press count distribution and on press 5, caused a rightward shift. **e)** Representative examples of a mouse's press counts from non-consecutive, selected trials with no stimulation, or with direct pathway excitation triggered on press 2 or press 5. **f)** For all mice, direct pathway excitation on press 2 did not affect the press count distribution and on press 5, caused a rightward shift. **g)** Representative examples of a mouse's press counts from non-consecutive, selected trials with no stimulation, or with indirect pathway excitation triggered on press 2 or press 5. **h)** For all mice, indirect pathway excitation on press 2 caused a leftward shift and narrowing of the press count distribution to counts of 2 and on press 5, caused a narrowing of the distribution to counts of 5. Bars represent the % of all trials across all mice.

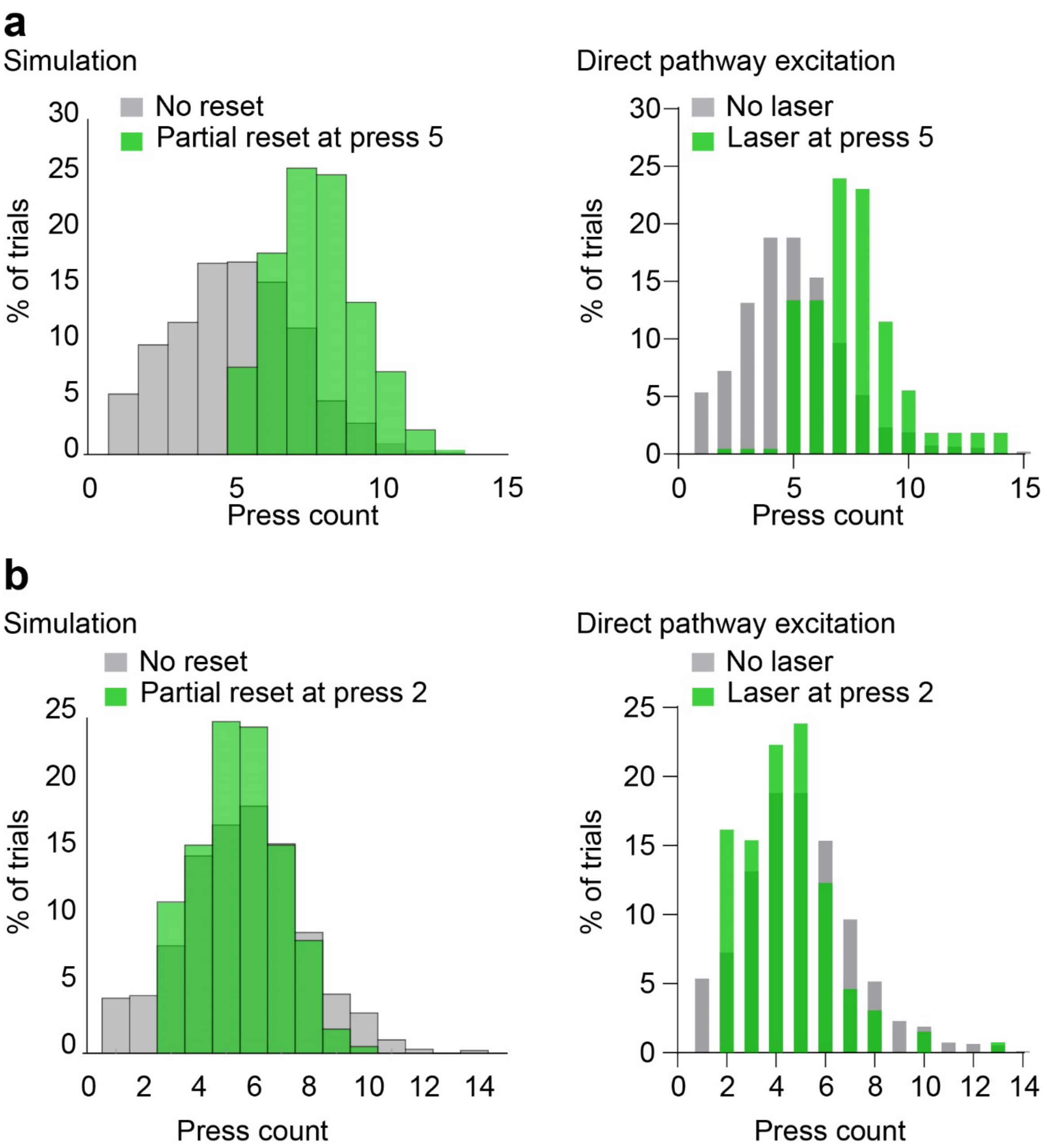


**Extended Data Fig. 6. Simulation of a partial count reset fits the empirical data. a)** Simulation of count estimation incorporating a partial reset (0.5) in the internal counting process on press 5. Direct pathway activation on press 5 caused mice to press as if their internal counter was partially reset. **b)** Simulation of count estimation incorporating a partial reset (0.5) in the internal counting process on press 2. Direct pathway activation on press 2 caused mice to press as if their internal counter was partially reset.

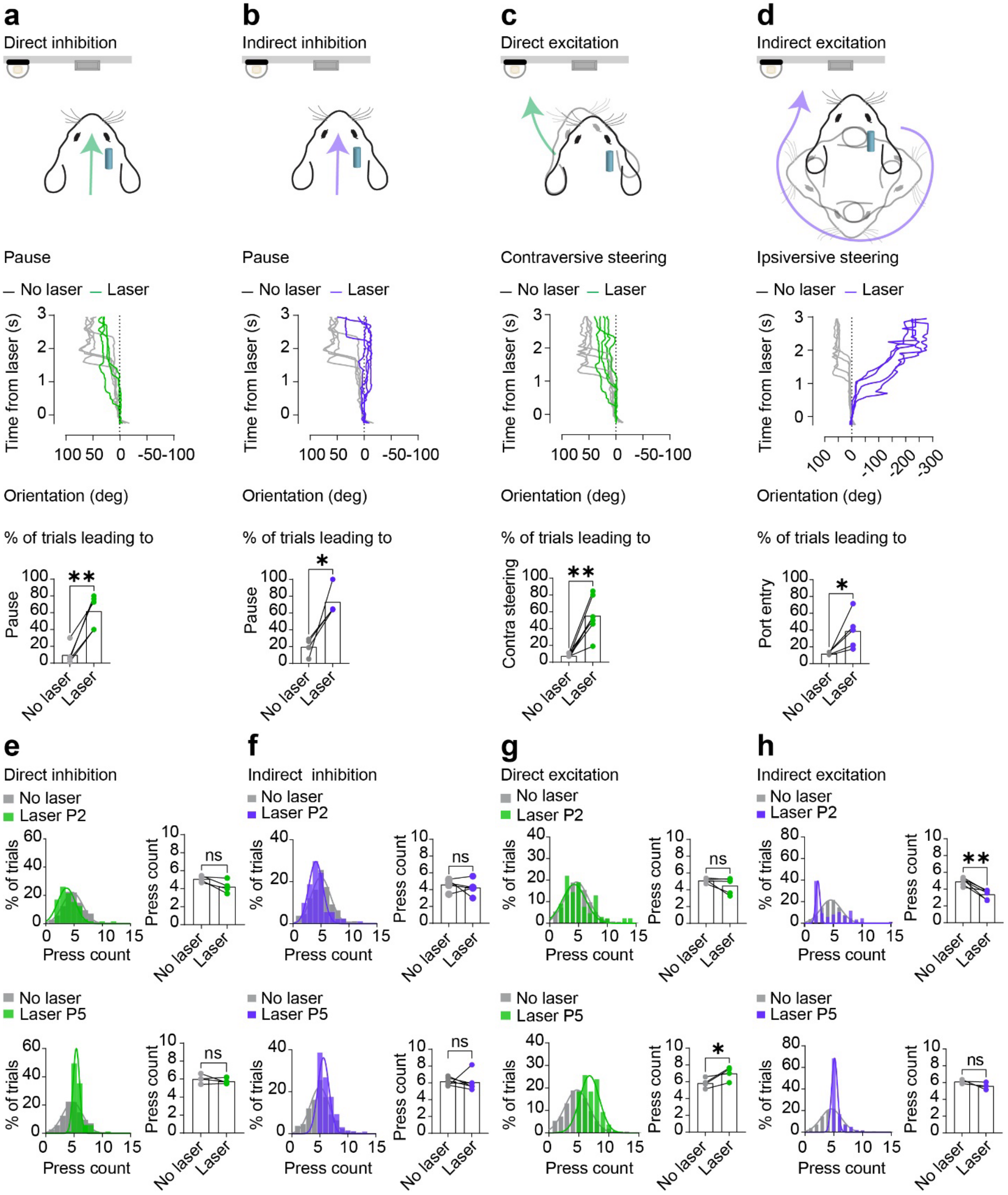


**Extended Data Fig. 7. Striatal effects on steering and press counting are hemisphere dependent. a-d)** Unilateral inhibition (500ms, 3-5mW) or excitation (500ms, 20-30Hz, 5-7mW) of the hemisphere contralateral to the port was triggered by press 2. **a)** Direct pathway inhibition caused a brief pause, the percentage of trials that led to a pause were significantly higher compared to eYFP controls (Paired t-test, $p$ = 0.018). **b)** Indirect pathway inhibition caused a brief pause, the percentage of trials that led to a pause were significantly higher compared to eYFP controls (Paired t-test, $p = 0.031$). **c)** Direct pathway excitation caused mice to steer contraversively towards the port, the percentage of trials that led to contraversive steering were significantly higher compared to no laser controls (Paired t-test, $p = 0.001$). **d)** Indirect pathway excitation caused mice to steer ipsiversively, all the way around to the port (360-degree turn). The percentage of trials that led to a port entry was significantly higher compared to no laser controls (Paired t-test, $p = 0.045$). **e-f)** Unilateral inhibition (500ms, 3-5mW) or excitation (500ms, 20-30Hz, 5-7mW) of the hemisphere contralateral to the port was triggered by the 2$^{nd}$ or 5$^{th}$ press. **e)** Direct pathway inhibition had no effect on the total number of presses in the stimulation trial or the press count distribution when compared to no laser controls (Paired t-test, P2: $p = 0.4583$, P5: $p = 0.393$). **f)** Indirect pathway inhibition had no effect on the total number of presses in the stimulation trial or the press count distribution when compared to eYFP controls (Paired t-test, P2: $p = 0.496$, P5: $p = 0.882$). **g)** Direct pathway excitation on press 2 had no effect on the total number of presses in the stimulation trial or the press count distribution when compared to no laser controls (Paired t-test, $p = 0.193$). When excitation occurred on Press 5, the number of presses in the stimulation trial were significantly greater than no laser controls (Paired t-test, $p = 0.043$). This resulted in a rightward shift in the press count distribution. **h)** Indirect pathway excitation on press 2 caused the number of presses in the stimulation trial to be significantly less when compared to no laser controls (Paired t-test, $p = 0.002$). This caused a leftward shift in the press count distribution. When excitation occurred on press 5, there was no effect on the total number of presses in the stimulation trial when compared to no laser controls (Paired t-test, $p = 0.147$). Bars represent the mean. * $p < 0.05$, ** $p < 0.01$.

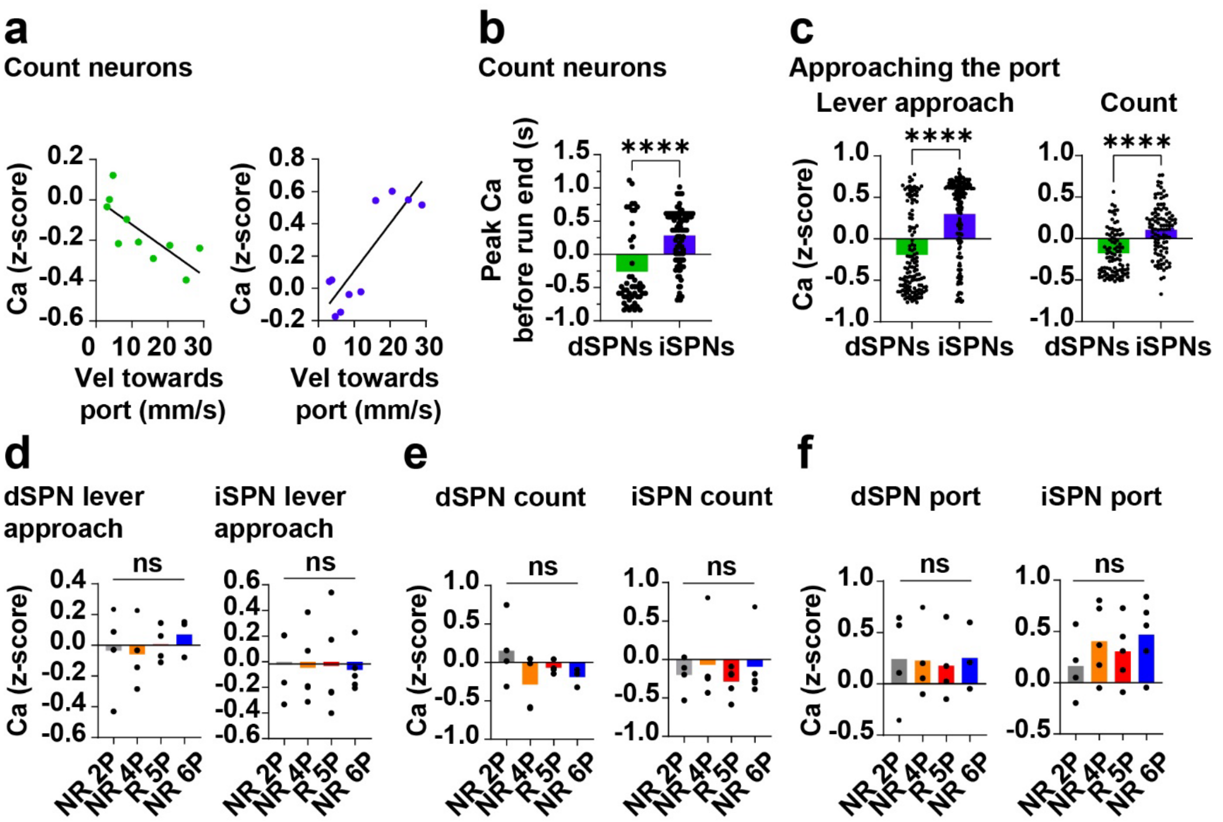


**Extended Data Fig. 8. dSPNs and iSPNs during counting and goal-directed approach behavior. a)** dSPN count neuron activity was negatively correlated with velocity to the port ($R^2 = 0.58$, $p = 0.01$) and iSPN count neuron population activity was positively correlated with velocity to the port ($R^2 = 0.73$, $p = 0.001$). **b)** For count neurons, dSPN activity peaked significantly before iSPN activity (Unpaired t-test $p < 0.0001$). **c)** For lever approach neurons, iSPN activity was significantly higher than dSPN activity when mice approached the port (Unpaired t-test, $p < 0.0001$). For count neurons, iSPN activity was significantly higher than dSPN activity when mice approached the port (Unpaired t-test, $p < 0.0001$). **d-f)** The activity of each SPN population on reward (R) and no-reward (NR) trials following different amounts of presses (e.g., NR 2P = no-reward 2 presses). Data represents mean population activity from 0 to 0.5s after port entry across mice. **d)** The activity of lever approach dSPNs was not significantly modulated by reward (RM ANOVA, $p = 0.81$). **e)** The activity of lever approach dSPNs and iSPNs was not significantly modulated by reward (dSPNs: RM ANOVA, $p = 0.13$, iSPNs: RM ANOVA, $p = 0.96$). **f)** The activity of count dSPNs and iSPNs was not significantly modulated by reward (dSPNs: RM ANOVA, $p = 0.17$, iSPNs: RM ANOVA, $p = 0.44$). Data represents mean ± SEM.

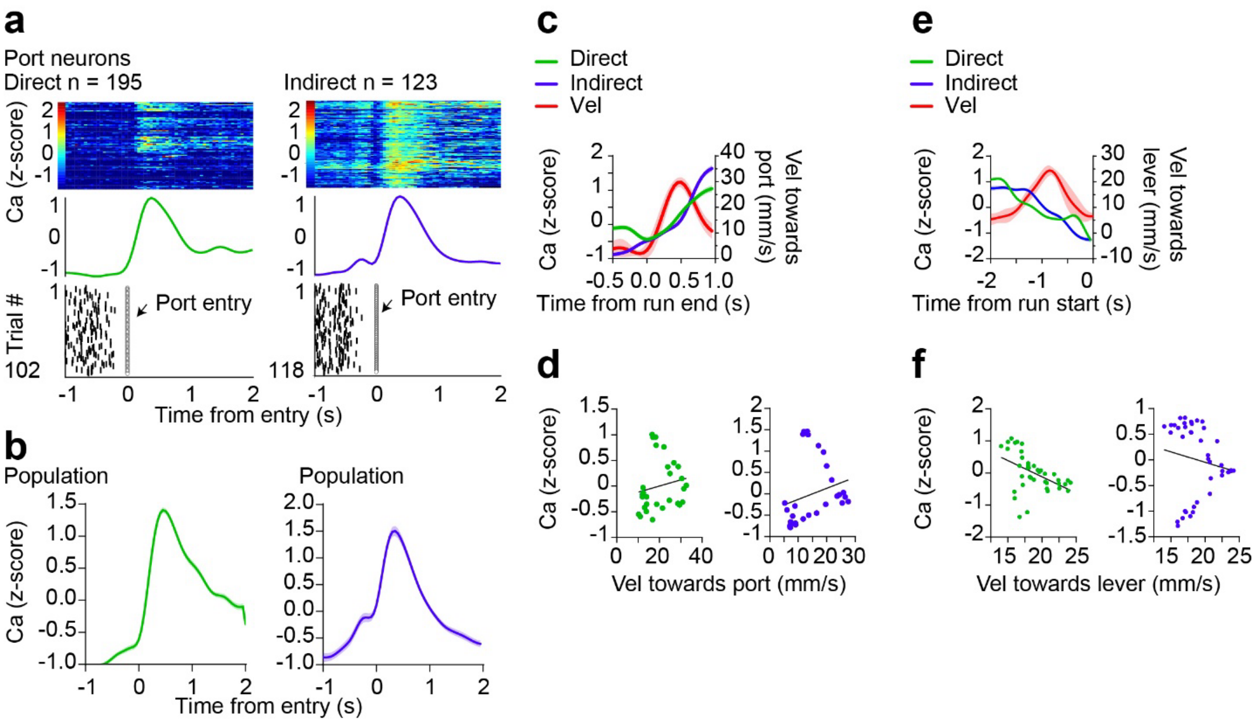


**Extended Data Fig. 9. Striatal neurons related to port entry do not represent left-right velocity. a)** Representative examples of dSPN and iSPN port neurons aligned to entry. **b)** The population of port entry neurons increased after mice entered the reward port. c) dSPN and iSPN port neuron population activity and port-approach velocity aligned to run end (transition to port). **d)** dSPNs and iSPNs were not correlated with velocity towards the port (dSPNs: $R^2$ = 0.03, p = 0.34; iSPNs: $R^2$ = 0.06, p = 0.17). **e)** dSPN and iSPN port neuron population activity and lever-approach velocity aligned to run start (transition to lever). **f)** dSPNs were negatively correlated with velocity towards the lever and iSPNs were not correlated with velocity towards the lever (dSPNs: $R^2$ = 0.22, p = 0.002; iSPNs: $R^2$ = 0.02, p = 0.33). Data represents mean ± SEM.

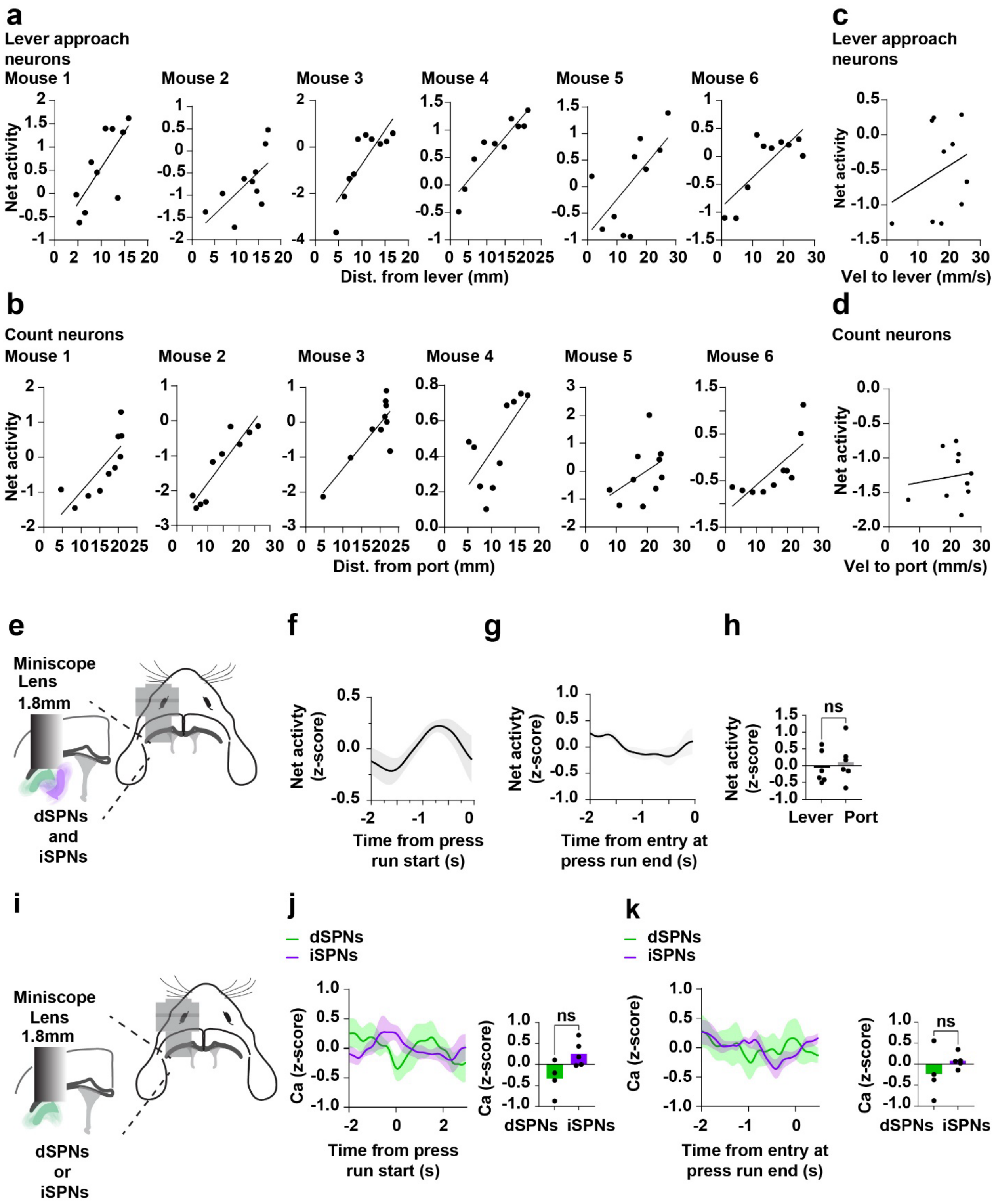


**Extended Data Fig. 10. Net activity (difference between dSPN and iSPN) was positively correlated with distance to goal targets. a)** The net activity of lever approach SPNs was higher when mice were further from the lever (mouse1: $R^2 = 0.54$, $p = 0.01$; mouse2: $R^2 = 0.43$, $p = 0.03$; mouse3: $R^2 = 0.69$, $p = 0.002$; mouse4: $R^2 = 0.42$, $p = 0.03$, mouse5: $R^2 = 0.84$, $p =$

0.0002, mouse6: $R^2 = 0.60$, $p = 0.008$). **b)** The net activity of port approach SPNs was higher when mice were closer to the port (mouse1: $R^2 = 0.62$, $p = 0.006$; mouse2: $R^2 = 0.83$, $p = 0.0002$; mouse3: $R^2 = 0.69$, $p = 0.002$; mouse4: $R^2 = 0.18$, $p = 0.2$; mouse5: $R^2 = 0.47$, $p < 0.02$, $R^2 = 0.56$, $p < 0.012$). **c)** Net lever approach activity was not correlated with velocity to the lever ($R^2 = 0.09$, $p = 0.39$). **d)** Net count activity was not correlated with velocity to the port ($R^2 = 0.018$, $p = 0.70$). **e)** To understand total net population dynamics, the average calcium activity of all dSPNs and iSPNs from each mouse in the two-color imaging experiments was taken. **f)** The net activity of all SPNs before press run start. **g)** The net activity of all SPNs before entry at press run end. **h)** The net activity at the lever was not significantly different than the net activity at the port (Paired t-test, $p = 0.59$). Data represents mean ± SEM. **i)** To understand total population dynamics, the average calcium activity of all dSPNs and iSPNs from each mouse in the single-color imaging experiments was taken. **j)** Total dSPN and iSPN activity as mice approached the lever. dSPN and iSPN activity was not significantly different as mice approached the lever (Unpaired t-test, $p = 0.056$). **k)** Total dSPN and iSPN activity as mice approached the port. dSPN and iSPN activity was not significantly different as mice approached the port (Unpaired t-test, $p = 0.37$).

**Video 1. Inhibition of dSPNs in the ipsilateral hemisphere on press 2.**

This video shows a representative example of unilateral inhibition of dSPNs (D1::GtACR) in the hemisphere ipsilateral to the reward port, triggered on the second lever press.

**Video 2. Inhibition of iSPNs in the ipsilateral hemisphere on press 2.**

This video shows a representative example of unilateral inhibition of iSPNs (A2a::GtACR) in the hemisphere ipsilateral to the reward port, triggered on the second lever press.

**Video 3. Excitation of dSPNs in the ipsilateral hemisphere on press 2.**

This video shows a representative example of unilateral excitation of dSPNs (D1::ChR2) in the hemisphere ipsilateral to the reward port, triggered on the second lever press.

**Video 4. Excitation of iSPNs in the ipsilateral hemisphere on press 2.**

This video shows a representative example of unilateral excitation of iSPNs (A2a::ChR2) in the hemisphere ipsilateral to the reward port, triggered on the second lever press.

**Video 5. Inhibition of dSPNs in the ipsilateral hemisphere on press 5.**

This video shows a representative example of unilateral inhibition of dSPNs (D1::GtACR) in the hemisphere ipsilateral to the reward port, triggered on the fifth lever press.

**Video 6. Excitation of iSPNs in the ipsilateral hemisphere on press 5.**

This video shows a representative example of unilateral excitation of iSPNs (A2a::ChR2) in the hemisphere ipsilateral to the reward port, triggered on the fifth lever press.

**Video 7. Inhibition of iSPNs in the ipsilateral hemisphere on press 5.**

This video shows a representative example of unilateral inhibition of iSPNs (A2a::GtACR) in the hemisphere ipsilateral to the reward port, triggered on the fifth lever press.

**Video 8. Excitation of dSPNs in the ipsilateral hemisphere on press 5.**

This video shows a representative example of unilateral excitation of dSPNs (D1::ChR2) in the hemisphere ipsilateral to the reward port, triggered on the fifth lever press.

**Video 9. Inhibition of dSPNs in the contralateral hemisphere on press 2.**

This video shows a representative example of unilateral inhibition of dSPNs (D1::GtACR) in the hemisphere contralateral to the reward port, triggered on the second lever press.

**Video 10. Inhibition of dSPNs in the contralateral hemisphere on press 5.**

This video shows a representative example of unilateral inhibition of dSPNs (D1::GtACR) in the hemisphere contralateral to the reward port, triggered on the fifth lever press.

**Video 11. Inhibition of iSPNs in the contralateral hemisphere on press 2.**

This video shows a representative example of unilateral inhibition of iSPNs (A2a::GtACR) in the hemisphere contralateral to the reward port, triggered on the second lever press.

**Video 12. Inhibition of iSPNs in the contralateral hemisphere on press 5.**

This video shows a representative example of unilateral inhibition of iSPNs (A2a::GtACR) in the hemisphere contralateral to the reward port, triggered on the fifth lever press.

**Video 13. Inhibition of dSPNs in the contralateral hemisphere on press 2.**

This video shows a representative example of unilateral inhibition of dSPNs (D1::GtACR) in the hemisphere contralateral to the reward port, triggered on the second lever press.

**Video 14. Excitation of dSPNs in the contralateral hemisphere on press 5.**

This video shows a representative example of unilateral excitation of dSPNs (D1::ChR2) in the hemisphere contralateral to the reward port, triggered on the fifth lever press.

**Video 15. Excitation of iSPNs in the contralateral hemisphere on press 2.**

This video shows a representative example of unilateral excitation of iSPNs (A2a::ChR2) in the hemisphere contralateral to the reward port, triggered on the second lever press.

**Video 16. Excitation of iSPNs in the contralateral hemisphere on press 5.**

This video shows a representative example of unilateral excitation of iSPNs (A2a::ChR2) in the hemisphere contralateral to the reward port, triggered on the fifth lever press.
